# Mapping individual differences across brain network structure to function and behavior with connectome embedding

**DOI:** 10.1101/2021.01.13.426513

**Authors:** Gidon Levakov, Joshua Faskowitz, Galia Avidan, Olaf Sporns

## Abstract

The connectome, a comprehensive map of the brain’s anatomical connections, is often summarized as a matrix comprising all dyadic connections among pairs of brain regions. This representation cannot capture higher-order relations within the brain graph. Connectome embedding (CE) addresses this limitation by creating compact vectorized representations of brain nodes capturing their context in the global network topology. Here, nodes “context” is defined as random walks on the brain graph and as such, represents a generative model of diffusive communication around nodes. Applied to group-averaged structural connectivity, CE was previously shown to capture relations between inter-hemispheric homologous brain regions and uncover putative missing edges from the network reconstruction. Here we extend this framework to explore individual differences with a novel embedding alignment approach. We test this approach in two lifespan datasets (NKI: n=542; Cam-CAN: n=601) that include diffusion-weighted imaging, resting-state fMRI, demographics and behavioral measures. We demonstrate that modeling functional connectivity with CE substantially improves structural to functional connectivity mapping both at the group and subject level. Furthermore, age-related differences in this structure-function mapping are preserved and enhanced. Importantly, CE captures individual differences by out-of-sample prediction of age and intelligence. The resulting predictive accuracy was higher compared to using structural connectivity and functional connectivity. We attribute these findings to the capacity of the CE to incorporate aspects of both anatomy (the structural graph) and function (diffusive communication). Our novel approach allows mapping individual differences in the connectome through structure to function and behavior.

## 1. Introduction

Understanding the neural basis of behavior is a fundamental goal in neuroscience. Traditionally, it has been addressed by relating the structure and activation patterns of individual brain regions to behavioral phenotypes or cognitive tasks. However, most complex behaviors are mediated not by a single, localized brain area, but rather by the integrated contribution of multiple regions forming a coherent, distributed network (Sporns, 2011). The connectome, a comprehensive map of the brain’s anatomical connections can be described and studied using methods and tools from network science. Research has gained valuable insight as to how individual nodes or modules in the network are organized to support specialized functional cognitive systems (Medaglia et al., 2015; Mišić & Sporns, 2016). In the past, most studies have aimed to detect differences in brain-behavior relations between groups of individuals. Increasingly, the focus has shifted to studying individual variations in connectome topology and its association with individual differences in brain function and behavior (Abdelnour et al., 2018; Lin et al., 2020).

The connectome is represented as a matrix comprising all dyadic connections among pairs of brain regions. Accordingly, a natural description of a single brain region would be a vector in this matrix, incorporating all its direct pairwise connections (a connectivity “fingerprint”; Passingham et al., 2002). Such representation is limited as it fails to capture relations of higher-order within the network (Goyal & Ferrara, 2018). While multiple graph descriptive measures have been proposed to capture local or global network attributes (Rubinov & Sporns, 2010), most are limited to describing only a specific network feature. Connectome embedding (CE) is an alternative approach for finding compact vectorized representations of nodes that capture their local and global topological attributes (Rosenthal et al., 2018). This approach, drawing from the field of natural language processing, is based on the Word2Vec algorithm (Mikolov et al., 2013) in which words are embedded in a low dimensional space that preserves their context as found in a given corpus of text. These representations were shown to capture semantic relations among words by applying simple vector arithmetic. For example, the result of combining vec(“King”) - vec(“Man”) + vec(“Woman”) was closer to vec(“Queen”) than to any other word in terms of its cosine similarity. Inspired by an adaptation of this approach to graph embedding (Grover & Leskovec, 2016; Perozzi et al., 2014), CEs are created by capturing a node’s “context” defined by its neighbors in a sequence of random walks on the brain graph. The resulting vectors were shown to capture relations between inter-hemispheric homologous brain regions and uncover putative edges that were missing from the structural network reconstruction (Rosenthal et al., 2018). This ability of the CE approach to capture meaningful topological attributes suggests that it could be used to improve the mapping of structural to functional connectivity and ultimately to behavior.

It has long been argued in neuroscience that structure determines function (Kristan & Katz, 2006). Accordingly, the nature of the correspondence between structural and functional brain connectivity is one of the core questions in the study of connectomes (Honey, Thivierge, & Sporns, 2010; Suárez, Markello, Betzel, & Misic, 2020). The two are distinct but complementary measures of brain connectivity. Structural connectivity quantifies the physical connections among neuronal elements, while statistical dependence between their time course is measured in functional connectivity. Although functional connectivity depends on the structural backbone, the observed correlation between the two is only moderate (Honey et al., 2009; Suárez et al., 2020). This, in part, can be attributed to higher-order interactions that drive functional connectivity but are missing in dyadic structural connectivity (Adachi et al., 2012). Such higher-order relations are captured in CE and indeed they were shown to account for a larger portion of the observed variance in functional connectivity (Rosenthal et al., 2018). These previous attempts were conducted at the group-level and it is crucial to examine whether such mapping could also be applied at the individual subject level and whether it would preserve inter-subject variability in structure-function relation.

Large scale data sharing initiatives have enabled applications of predictive modeling to study individual differences and validate them within and across large cohorts. Nevertheless, employing such techniques to connectivity data remains challenging due to their high complexity and dimensionality. CE offers a promising complementary framework that addresses these challenges by representing nodes in low-dimensional space while preserving their context in the global network topology. However, utilizing these nodal representations for predicting function and behavior requires their mutual alignment to the same latent space to allow comparison across individuals. Multiple fitting iterations of the Word2Vec algorithm, even on the same brain graph, result in sets of vectors with similar relative positions among each other but with different absolute values (Dev et al., 2019; for the stability of the relations among embeddings see Wang et al., 2020). In the domain of machine translation, this is typically addressed by finding a transformation that minimizes word distances from one language to another (Smith et al., 2017). However, such an approach could not be easily adopted to align brain nodes across individuals as it also eliminates parts of the variability associated with differences in the underlining structural connectivity. Hence, applying the CE framework to explore individual differences requires an alignment method that preserves this structural variability.

In the current work, we advance the CE framework by presenting a novel approach which enables us to align separately learned embeddings to a common latent space (Fig. 1). The alignment is based on a closed-form solution that utilizes parameters identified during the embedding fitting process. We validate this approach using two large lifespan cohorts (Nooner et al., 2012; Taylor et al., 2015) that include diffusion-weighted imaging, resting-state fMRI, as well as demographics and cognitive performance measures. We align CE within and between individuals and test the alignment effect on the similarity of nodes and the relations between nodes across embeddings. We then demonstrate that CE can improve the mapping of structural to functional connectivity at the individual level as previously done at the group-level (Rosenthal et al., 2018). We further examine whether this mapping preserves age-related variance in structure-function correspondence at the network-level and at the edge-level. Interpreting the CE as a generative model of diffusive communication around nodes, we test the effect of the random walk parameters on structure-function mapping. Finally, we examine whether the aligned embeddings could be used to predict age and intelligence in a held-out sample. We include an in-depth derivation of the CE fitting and alignment process and share a set of interactive notebooks reproducing the main findings and a python package implementation (Cepy; https://github.com/gidlev/cepy).

**Fig. 1.**
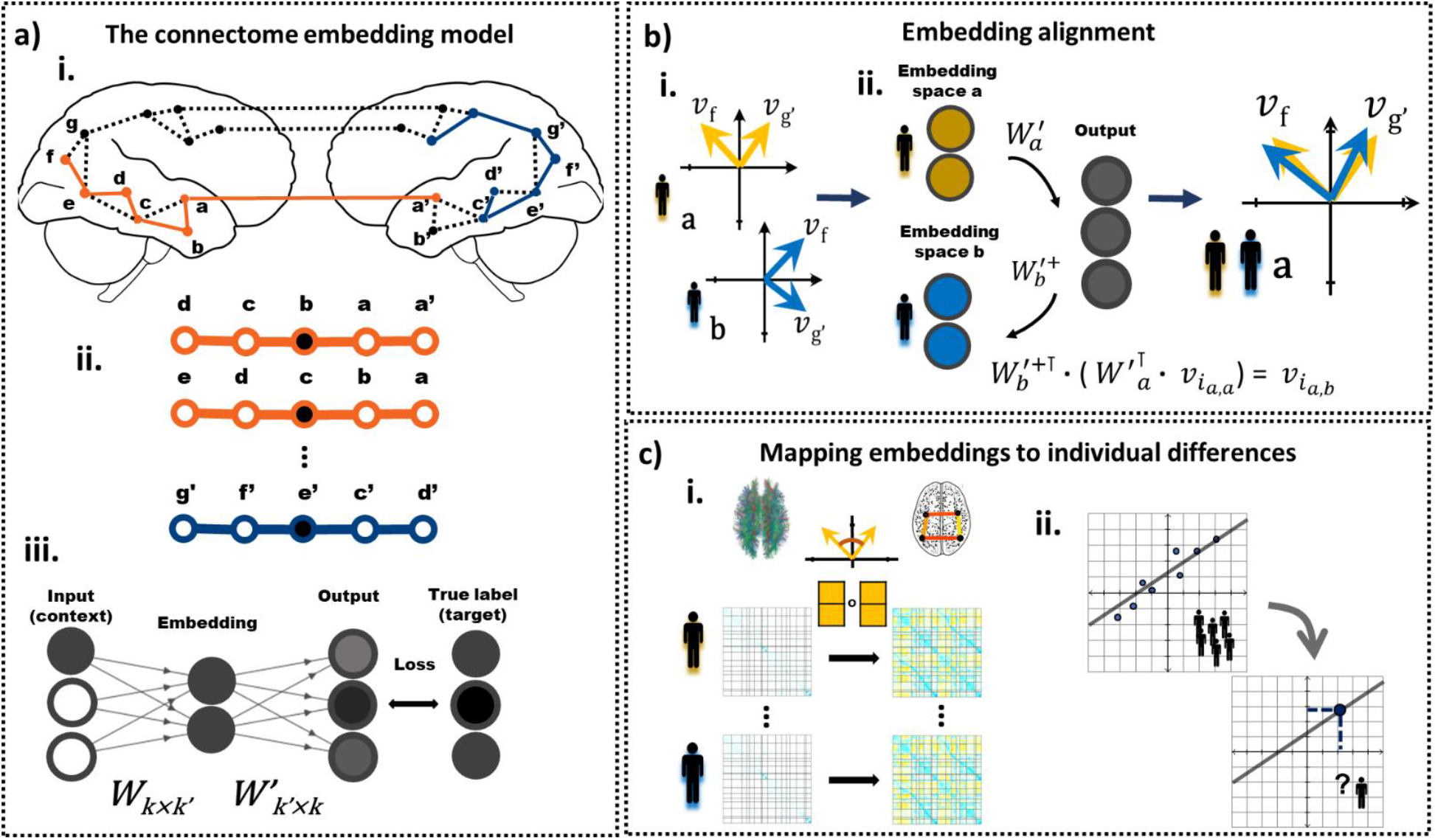
A general outline of the connectome embedding framework and its usage at the individual subject level. a.i. The structural connectome is the input to the CE model. Multiple random walks are sampled from the structural connectivity graph. Letters represent all unique nodes in both hemispheres, dash lines represent edges that exist in the connectome and colored line represent sampled random walks. a.ii. Sliding, fixed size windows are taken over random walk sequences. Within each window, the center node is used as the target (black) and the surroundings as context (white). a.iii. Pairs of context and target nodes are used as the input and target of an artificial neural network with a single hidden layer, i.e. the embedding layer. The input, output and target layers are *k* dimensional vectors, where *k* is the number of nodes in the brain graph. The embedding layer is a k’ dimensional vector, where *k’* is set to be *k’*<*k*. *W* and *W’* are the learned weight matrices that define the transformation between the input and embedding layer, and the embedding and output layer, respectively. The model parameters,*W* and *W’,* are iteratively updated using stochastic gradient descent. b.i. Independent fits of the model (e.g. in different subjects) results in embedding vectors with similar angle among node pairs but different absolute values. Embedding alignment is required to allow a comparison of different CE. b.ii. Here, embeddings are transformed from the latent space of subject a (yellow) to the latent of subject b (blue) to allow their comparison. This is done in two stages, first, we apply the *W’_a_* transformation from the embedding to the output space.Then, we apply the pseudo-inverse of the *W’_b_* transformation from the output back to the embedding space. In the embedding vectors the first subscript *i* denotes the node’s index and the second the source and current subject’s latent space. Similarly, the subscript in the transformation matrices denotes their associated subject. The same approach could be similarly used to align independent embedding iterations of the same subject. c.i. The pair-wise cosine similarities or the element-wise multiplication among embedding vectors are used to map structural to functional connectivity. c.ii. The aligned embeddings vectors are used for out-of-sample prediction of individual differences.

## 2. Materials and Methods

### 2.1 Participants

The data were taken from two lifespan large-scale cross-sectional studies that included functional, structural, and diffusion brain magnetic resonance imaging (MRI) along with demographics and behavioral data. The first dataset is the enhanced Nathan Kline Institute-Rockland Sample (eNKI-RS; Nooner et al., 2012) and the second is the Cambridge Centre for Ageing and Neuroscience dataset (Cam-CAN; Shafto et al., 2014) referred here as dataset 1 (DS1) and dataset 2 (DS2), respectively. DS1 is composed of 542 subjects (305 females, 192 males) aged 7-84 recruited from Rockland County, USA. All participants provided informed consent and the study was approved by the Institutional Review Board at the Nathan Kline Institute (#226781 and #239708) and Montclair State University (#000983 A and #000983B). The data is openly available online at http://fcon_1000.projects.nitrc.org/indi/enhanced/neurodata.html. DS2 includes 601 subjects aged 18-87 roughly uniformly distributed from Cambridge City, UK. All participants provided informed consent and the study was approved by the local ethics committee, Cambridgeshire 2 Research Ethics Committee (reference: 10/H0308/50). The data is freely available upon online access request https://camcan-archive.mrc-cbu.cam.ac.uk/dataaccess/. Additional information on the recruitment, eligibility criteria and demographics of both samples are available in the relevant publications (Nooner et al., 2012; Shafto et al., 2014). Exclusion criteria included successful completion of the preprocessing and quality control stages and are specified in the methods (2.3) and in the supplementary information (SI 2).

### 2.2 MRI acquisition

DS1 and DS2 were both acquired on 3T Siemens scanners. For each subject of each dataset, a T1-weighted image (T1w), diffusion-weighted image (DWI), and resting-state functional MRI (rsfMRI) were acquired. Both datasets include standard T1w anatomical acquisitions. DS1 includes DWI with 128 directions in one shell with a b-value of 1500 s/mm^2^, and DS2 includes DWI with 60 directions over two shells of 1000 and 2000 s/mm^2^. DS1 includes a rsfMRI acquisition with a relatively short repetition time of 0.645 seconds with a scan time of 9.68 minutes, and DS2 includes a more standard acquisition scheme with a repetition time of approximately 2 seconds with a scan time of 8.56 minutes. The specific parameters of each acquisition in each dataset can be found in SI table 1.

### 2.3 MRI preprocessing

Here we report a general overview of the independent preprocessing pipelines applied for DS1 and DS2. Given recent reports of the effect of processing strategies on MRI analysis (Botvinik-Nezer et al., 2020), we intend to show converging results across two datasets employing different processing pipelines. Complete preprocessing details for both datasets are included in SI Section 1.

T1w scans from both datasets were preprocessed through FreeSurfer’s (version 6.0) recon-all processing stream. FreeSurfer’s cortical segmentation and spherical warp were used to transfer parcellations to each subject’s volumetric anatomical space. In the case of DS1, the Schaefer 200-node cortical parcellation was rendered (Schaefer et al., 2018), where as, in DS2, the Connectome Mapping toolkit was used to render the Lausanne 233 node parcellation (219 cortical, 14 subcortical; Gerhard et al., 2011).

DWI of both datasets were preprocessed with pipelines that included the following steps: denoising, motion and eddy current distortion correction, and alignment to the T1w using FreeSurfer’s white matter segmentation (Ades-Aron et al., 2018; Bathelt et al., 2017). For both datasets, local orientation modeling and tractography was run via the Dipy package (Garyfallidis et al., 2014). Constrained spherical devolution was used to fit a local orientation model at each voxel, with a spherical harmonic order of 8 in DS1 and 6 in DS2. For DS1, probabilistic streamline tractography was conducted after seeding each white matter voxel five times. For DS2, deterministic streamline tractography was conducted with a seeding density of 27. In both datasets, streamlines shorter than 10mm or ones that did not terminate in grey matter were discarded.

Functional images of DS1 were preprocessed with fMRIPrep (version 1.1.8; Esteban et al., 2019) and images of DS2 were preprocessed with the Configurable Pipeline for the Analysis of Connectomes (C-PAC; Cameron et al., 2013). Briefly, both pipelines included the following steps: slice-timing correction, motion correction, skull stripping, and estimation of motion parameters and other nuisance signal time series. For DS1, preprocessed images were rendered in the subject’s T1w space, at the original resolution of the rsfMRI. For DS2, preprocessed images were rendered in rsfMRI space.

For DS1 and DS2, structural connectivity matrices were constructed by counting the number of streamlines between regions normalized by the volume of these regions, rendering a streamline density. For DS1 and DS2, subjects for which more than 10 nodes in the structural connectivity matrix had a degree of 0 were excluded and were not used in subsequent analysis (DS1: none omitted, 542 left; DS2: 6% omitted, 601 left). The samples of both datasets were divided into a training (67%) and test set (33%) for subsequent analyses that required out-of-sample accuracy estimation (DS1: 361 and 181, DS2: 401 and 200; training and test subjects, respectively).

For DS1 and DS2, functional connectivity matrices were rendered after filtering the functional volumes for nuisance signals. For DS1 rsfMRI, the first four frames were dropped. These data were then bandpass filtered (0.008 – 0.08Hz) and confound regressed in a manner orthogonal to the temporal filters using 6 motion estimates, the mean time series derived in CSF, WM, and whole brain masks, the derivatives of these nine regressors, and the squares of these 18 terms. Spike regressors were added for each frame with framewise displacement above 0.5mm. Data were linearly detrended and standardized. Exclusion criteria included greater than 15% spike frames and outlier image quality metrics (4% omitted; 542 subjects left; for more details, see SI 6.2). For DS2 rsfMRI, regression of the first 5 principal components of signal from white matter and CSF (Behzadi et al., 2007), 6 motion parameters and linear and quadratic trends, global signal regression, followed by temporal filtering between 0.1 and 0.01 Hz and. Finally, a scrubbing threshold of 0.5mm frame-wise displacement was applied (Power et al., 2014; removal of 1 TR before and 2 TR after excessive movement). Exclusion criterion for excessive movements was determined a priori to less than 50% (4 min and 20 sec) of the resting-state session after the scrubbing procedure (25% omitted; 452 subjects left). For DS1 and DS2, functional connectivity as used in all analyses, was defined as the Pearson correlation among pairs of ROIs’ time series followed by Fisher’s r-to-z transformation.

### 2.4 Intelligence assessment

Structural connectivity was previously associated with behavioral measures of intelligence (Booth et al., 2013; Penke et al., 2012), and here was used to test the ability of the CE approach to capture individual differences in behavior. In DS1 general intelligence was assessed using the full scale of the Wechsler Abbreviated Scale of Intelligence (WASI-II; Wechsler, 1999). Subjects in DS2 underwent the Cattell Culture Fair test, Scale 2 Form A that aims to measure fluid intelligence independently of cultural differences (CFIT; Cattell & Cattell, 1973). In the relevant analyses, we used a sample of subjects in training and test sets for whom we had the structural connectivity matrices and intelligence behavioral scores (DS1: 528; d2: 587). Next, to remove outliers, participants with intelligence scores larger than 2 standard deviations from the mean were omitted from the analyses (DS1: 0% omitted, 528 left; DS2: 4% omitted, 565 left). The WASI-II full scale in DS1 produced age-adjusted scores that were uncorrelated with age within the sample (r(526)=.038, p=.386). CFIT appeared in DS2 as the raw accuracy score for all test items. This score was significantly correlated with age (r(563)=-0.652, p<2.2e-16) and hence constitutes an age-related measure of intelligence.

### 2.5 Node embedding -general outline and the random walk sampling

As in previous work (Rosenthal et al., 2018), we used the word2vec algorithm (Mikolov et al., 2013) to create a vectorized representation of brain nodes based on their high-level topological relations. Originally, the word2vec algorithm was used to create word embeddings that preserve their context as it typically appears in a sentence. Specifically, the model is given a corpus of words *w* and their context *c*. Its goal is to learn a set of parameters θ by maximizing the conditional probability p(c|w; θ) for the skip-gram model or p(w|c; θ) for the continuous bag of words (CBOW) model (Goldberg & Levy, 2014). Simply put, the model’s goal is to predict the context given the target word or a target word given its context. The former is used in the current work. Recently, word2vec has been applied for embedding graph nodes instead of text (node2vec: Grover & Leskovec, 2016; Deepwalk: Perozzi, Al-Rfou, & Skiena, 2014). Here, for embedding brain graphs, we used a sliding, fixed-size window *s* taken from a sequence of a parameterized random walk on the brain graph (*s*=3). In each training sample, the center node in the sequence is the target *w* and the surrounding nodes are the context *c*. The surrounding nodes are defined as the *s* nodes that appeared before and *s* nodes that appeared after the target node *w* within the walk sequence. Training samples were produced by initiating *o* parameterized random walk sequences of length *l* from each node (*o* = 800, *l* = 20). We further elaborate on the parameterized random walk procedure in section 2.10. All the model’s parameters were set to be identical to those used by Rosenthal et al. (2018).

### 2.6 Node embedding – model implementation and parameters estimation

In its simplest form, the model represents a fully connected artificial neural network (ANN) with one hidden layer, i.e. the embedding layer, with no activation function. Both the context {c_1_,…,c_2×s_} and target *w* nodes are represented using a “one-hot encoding”, meaning that node *i* is encoded as a vector with zero in all its entries except the *i*^th^ position that is equal to one. The number of neurons in the input and output layers is the number of nodes in the graph *k* and the number of neurons in the embedding layer is set to *k’*, when typically, *k’*<*k* (here *k’* = 30). The transformation between the input, embedding and output layers, denoted here as *v^input^, v^embedd^*, *v^output^*, are defined using two weight matrices. A *k* × *k’* matrix *W* between the input and the embedding layer, and a *k'* × *k* matrix *W’* between the embedding layer and the output layer:

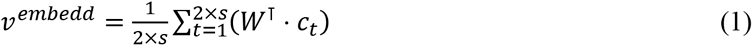

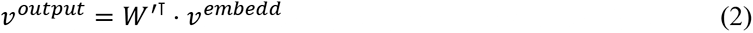

Notice that the embedding layer is computed as the average of all vector-matrix product of the context node vectors with the *W* matrix. Both *W* and *W’* are learned by fitting the model on a given sample of random walks. Effectively the model performs a classification task in which the input is the context nodes {c_1_,…,c_2×s_} and the goal is to predict the target node *w*. This is done by first applying a Softmax function to the output layer which normalizes its entries into a probability distribution that sums to one. Here for the *i*^th^ entry:

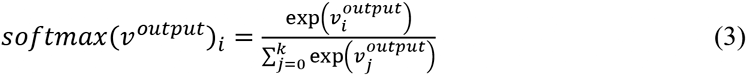

Next, the model’s loss is computed by taking the negative log, i.e. the logarithmic loss, of the target node *w* entry, its index marked here as *:

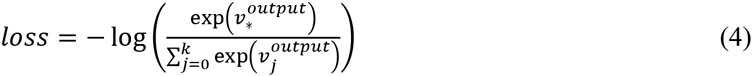

Here in simplified form:

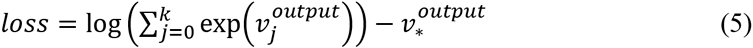

Finally, for a given training sample, we update the model’s parameters θ, i.e. the weight matrices W and W’, by taking the derivative of the loss with respect to each matrix. The parameters are iteratively updated after each observation of a single sample or a batch of training samples, i.e. using stochastic gradient descent. For a more in-depth review of the model and training procedure including implementational details that are beyond the scope of the current work such as negative sampling, see Rong (2014).

### 2.7 CE alignment

Independent fitting iterations of the node2vec algorithm resulted in sets of vectors with similar cosine angle between each node pairs, but not necessarily similar absolute values (Dev et al., 2019). Here we demonstrate our novel approach which enables us to align separately learned CE to the same latent space (see Fig. 1.b.). As outlined in section 2.5, the ANN model includes two distinct representations. The first, which appears in the input and output layers, is defined by the one-hot encoding in which each entry corresponds to a particular node (see section 2.6). The second is a latent representation, the representation of the embedding layer, that is unique to each trained model. Notice that the learned matrices *W* and *W’* encode the transformation between these two representations and for this reason, we refer to them here as *transformations*. The first transformation *W* reduces the input dimensions from *k* to *k’* and effectively contains the embedding of each node. The second transformation *W’* increases the dimensions of the embedding layer *k’* again to the number of dimensions in the output layer *k*. While typically after fitting the algorithm on the data, only the first matrix *W* is retained, we suggest utilizing the second matrix *W’* as a transformation from one latent space to another. Specifically, given two separately trained models with different latent embedding spaces *a* and *b,* we want to transform the latent representation of node *i* from space *a* to *b*. In each of the models, a separate *W’* transformation was learned, *W’*_a_ and *W’*_b_, respectively. First, the embedding of node *i* in space *a* could be derived by taking the *i^th^* row of *W*_a_, or equivalently multiply the one-hot encoding of node *i*, *v^input^*_*i*_, with *W*_a_:

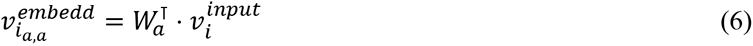

Note that for the embedding and output layers the first subscript denotes the node’s index and the second the source and current latent or output space. Next, the vector is multiplied with *W’*_a_ to transform it to the ANN output representation:

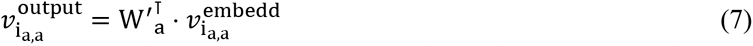

We assume that since the one-hot representational constraint exists in all fitting iterations, this representation is similar across separately trained models. Stemming from this, we can apply the inverse of transformation *W’*_b_, to move to latent space *b*. Since W’ is non-square we apply the pseudo-inverse transformation *W’*_b_^+^:

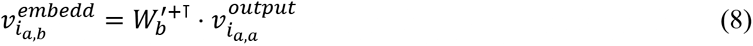

(7) and (8) could be summarized to a single step, incorporating the transformation of the embedding of node *i* from latent space *a* to latent space *b*:

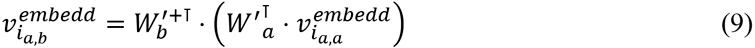

This alignment method does not require any additional learning or optimization process and relies on the transformation already learned within the ANN. This method could be used to align embeddings originated from separate fitting iterations. These could be fitted on the same structural connectivity matrices, from matrices taken from different individuals, or matrices of the same individual across time. As an additional step to improve embedding stability over fitting iterations (Wang et al., 2020), in the subsequent analysis we averaged multiple embeddings of the same individual after aligning them to the same space (100 times, unless stated otherwise). Python notebooks are available online to demonstrate the random walk sampling, word2vec model, model fitting, and the alignment process – https://github.com/gidlev/cepy/tree/master/examples.

### 2.8 Intra-individual CE similarity evaluation

We examine the effect of embedding alignment on the similarity of identical nodes across separate embedding iterations. For a random sample of subjects (n = 100) we ran the node2vec algorithm multiple times (m = 50), then aligned the resulting vectors to a common space and averaged them. The common reference space was created by applying the node2vec algorithm on the group consensus structural connectivity matrix. This procedure was repeated twice and the node’s embedding similarity across the two learned CE was quantified using cosine similarity. The similarity measure was averaged across all nodes (DS1: 200, DS2: 233) and calculated for all subjects. Next, we evaluated the similarity of the cosine angle among pairs of nodes across CEs that resulted from independent fitting iterations. This was done by calculating the displacement vector, i.e. the result of vector subtraction, among all possible pairs of nodes (DS1: 19900, DS2: 27028). Similarly, the cosine angle among the displacement vector obtained from the two separate learned CE was taken. The similarity measure was averaged across all node pairs and calculated for all subjects. In both tests, the similarity measures were compared to a case where the alignment step was omitted (“non-aligned” condition).

### 2.9 Inter-individual CE similarity evaluation

To test the quality of the alignment, we applied the embedding ranking test (Rosenthal et al., 2018). First, multiple node2vec iterations (m = 100) were fitted to each subject, the resulting embeddings were aligned to a common space (see section 2.7) and averaged. Then, for every node in one individual, we obtained its cosine similarity to all nodes in a second individual. A ranking test score of *k* meant that for a particular node, its corresponding node in the second individual was the k-closest node. The test was conducted for each node and every possible subject pair (n = 100; 9,950 pairs). Finally, we tested the similarity of relations among pairs of nodes across individuals. This was done using an analogy ranking test. The test measured whether the displacement vector from node *a* to *b* in one individual would express the same relation in another individual. Here, the ranking score was conducted by adding the displacement vector from node *a* to *b* taken from one individual to node *a* of another individual. The query, in this case, was how close is the resulting vector to node *b* of the second individual. This test was repeated for every subject and node pairs resulting in *k*-times more computations (DS1: k=200, DS2: k=233). Thus, due to computational time considerations, the analysis was conducted on a smaller, randomly selected, set of subjects (n = 30). Both tests were compared to a condition in which the alignment step was omitted.

### 2.10 Group-level structure to function mapping

Analyses on the group-level structure to function mapping were conducted on the consensus structural and functional connectivity matrices derived from all subjects in the test set. The functional group connectivity matrix was calculated by averaging the Fisher-r-to-z transformed correlations across all subjects. The structural group consensus connectivity matrix in each dataset was derived by averaging over all edge values that were present in at least 25% of subjects (de Reus & van den Heuvel, 2013). Structural edges on all other node pairs were assumed to be absent and set to zero in the consensus average. The group-level CE was created by applying the node2vec algorithm to the structural group consensus matrix. Structure-function correlation was quantified as Spearman’s correlation between all unique direct edges, i.e. edges that exist in the consensus matrix, for the structural connectivity. The same correlation measure was used for the CE cosine edges but applied to all edges, both direct and indirect. We used Spearman’s rank correlation because of the exponential distribution of the structural connectivity values. This measure was used in all subsequent connectome-level structure-function correlation assessments. To examine the contribution of individual edges to the increased CE-based structure-function mapping, we adopted the Leave-One-Trial-Out scheme (Gluth & Meiran, 2019). This was done by calculating Spearman’s rank correlation between the CE cosine similarity edges and their corresponding functional edges. This was followed by computing the difference in this correlation after removal of each of the edges. The correlation before minus after the removal, or Δ correlation, could inform us about the effect of a given edge on the overall correlation by examining its sign and relative magnitude.

### 2.11 Parameters of the random walk and their relation to structure to function mapping

In a fully diffusive communication process, signals propagate on the network structure driven only by local connectivity properties (Avena-Koenigsberger et al., 2018). In the node2vec algorithm, random walks are biased and assume some information about the node’s local neighborhoods. Two parameters guided the biased random walk, the return parameter *p* and the in-out parameter *q* (Grover & Leskovec, 2016). Specifically, the parameter *p* sets the likelihood for a random walker to immediately revisit a node, i.e. that at time *t*+*1* it would occupy the same node visited at time *t-1*. The parameter *q* controls the likelihood of a random walker to visit nodes that are not directly linked to the node visited in the previous step, i.e. that its step in time *t*+*1* would be to a node with edge = 0 with the node visited in time *t-1*. While high *p* and low *q* would guide the walk to remain in the vicinity of the initial starting point (local bias), the opposite would promote exploration of distant nodes (global bias). In all analyses reported here, we used a locally biased random walk (*p*=0.1, *q*=1.6; Rosenthal et al., 2018). Additionally, we examined the effect of manipulating *p* and *q* on the observed correlation between the CE cosine similarity matrix and the functional connectivity matrix. We report this correlation for randomly selected subjects from the training set (n=25) with functional and structural connectivity measures. We report this correlation separately for direct and indirect edges. Parameterized random walks were produced using the reference node2vec Python implementation (https://github.com/aditya-grover/node2vec).

### 2.12 Predicting group-level functional from structural connectivity with deep learning

Previous work (Rosenthal et al., 2018) has shown that a CE-based structure to function mapping could be further improved using a deep neural network. Here we adopted the same methodology in which the element-wise multiplications of any pair of node embeddings were used as input and the functional connectivity between the two as the target label. We used a fully connected, 4-layer network with 256 neurons in each layer. The network was implemented in Keras (François Chollet and contributors, 2015) and Tensorflow (Abadi et al., 2016). The training was conducted using an Adam optimizer, a learning rate of 0.001, and 250 epochs. We used 3-fold cross-validation by randomly dividing the edges of the mean functional connectivity matrix into 3 folds. In each iteration, two folds were used for fitting and the remaining fold was used for prediction. We additionally applied cross-validation on the group matrices such that edges and CEs used for fitting were taken from subjects within the training set and CE used for the prediction was taken from subjects within the test set. Finally, we wanted to account for the fact that nodes from the same community, defined by the 7 canonical resting-state networks (Yeo et al., 2011), have similar functional connectivity patterns. For this reason, we divided the 3-folds such that each intersection of two communities would not be repeated across folds. For example, all edges connecting a visual node to a motor node appeared only in one of the folds. This scheme was used for the created CE-based predictions for all possible functional edges, both direct and indirect.

### 2.13 Individual-level structure to function mapping

CE could be further utilized to map structural to functional connectivity at the individual subject-level. The following analyses were conducted for individuals that underwent DWI and a resting-state session, and were not excluded at the preprocessing stage (see section 2.3; DS1: 307 and 145, DS2: 361 and 181; subjects left in the training and test set, respectively). Structure-function correspondence was evaluated using subjects within the test set both for structural connectivity values and for the CE cosine similarity measure. Differences in structure-function mapping between CE and the connectivity values were examined using a paired-sample t-test between the correlation coefficient values. Finally, a deep learning model for structure-function mapping was applied using the same architecture and hyper-parameters described in section 2.10. Cross-validation was conducted both at the edge and subjects levels. On the edge-level we used the same 3-fold strategy as described in section 2.12. At the subjects-level, model fitting was conducted only on within the training set and testing on test set. We reported structure-function connectivity correlation separately for direct edges, indirect edges, and all edges.

### 2.14 Age-related changes in individual-level structure-function correspondence

Age was previously found to be associated with a decrease in the strength of structure-function connectivity correlation (Zimmermann et al., 2016). Here we wanted to examine whether this negative correlation between age and structure-function relation is preserved when relating CE to functional connectivity. The correlation was calculated for all the subjects within the test set. Structure-function correspondence was evaluated using Spearman’s rank correlation only on direct edges. Comparing the magnitude of the aging effect on the structure-function correspondence was evaluated using Steiger’s t-test for comparing dependent correlation values (Steiger, 1980; python implementation: https://github.com/psinger/CorrelationStats/). To test how age relates to edgewise structure-function correspondence we used the Leave-One-Trial-Out scheme described in section 2.10. The derived contribution score that measures the effect of a single node on the overall structure-function relation was applied for the same group of subjects. We then measured the Pearson correlation of each edge with age across subjects. Finally, we adapted the network contingency analysis method (Sripada et al., 2014). This method examines whether sub-blocks derived from each of seven canonical resting state networks (Yeo et al., 2011) present a larger number of edges correlated with age above a predetermined threshold than expected by chance, with chance defined through permutation testing. We repeated our analysis for a wide range of thresholds (|r| > 0.25, 0.3, 0.35, 0.4). To establish the null distribution, we used 10,000 permutations of ages and only accepted cells for which none of the correlations obtained after permutation was larger than the empirical value (see SI 8 for a full description of the analysis). Finally, we examined whether within the 7 canonical functional networks, within or between hemispheres, the supra-threshold edges were significantly larger or smaller than zero. We used a one sample t-test were the null hypothesis was that the population mean is zero.

### 2.15 Modeling individual differences in age and intelligence using CE

To test the ability of the CE framework to predict individual differences we trained a linear model to predict age and intelligence. First, a separate model was trained for each node and each input type, i.e. the CE, the CE cosine similarity matrix, the structural or the functional connectivity values as input. We used the rows of the CE cosine, structural and functional connectivity matrices and the nodal CE representations as the node-level inputs. The performance measure of the model was compared to the one obtained by fitting the same model to the structural or functional connectivity values. We used linear regression to predict age and intelligence. The performance was quantified as the Pearson correlation between the observed and predicted age or intelligence. The connectome-level model was created by taking the mean of all nodal-level predictions or training a second-level ensemble on them. Due to the high multicollinearity of the first-level prediction, an L2 regularized linear model trained with SGD was used as the ensemble model. As in the first-level model, the ensemble was trained within the training set and tested on the test set. Both models were implemented in Scikit-learn (LinearRegression and SGDRegressor; Pedregosa Fabian et al., 2011) with the default parameters. We additionally examined whether our results are confounded by a possible effect of gender, for age prediction, and gender and age, for intelligence prediction. We controlled for the two variables using linear regression by predicting the desired outcome, age and intelligence, with the covariates, gender or age and gender, as predictors, keeping only the residual.

## 3. Results

### 3.1 Individual CE alignment evaluation

The Node2vec algorithm was previously shown to capture high-order structural relations within group-averaged representations of the human connectome (Rosenthal et al., 2018). Extending the approach to individual brain networks requires that individual connectome embeddings are mutually aligned to allow an assessment of individual differences. Fitting the CE algorithm on a given individual graph results in an embedded vector representation for each brain node. Multiple fitting iterations result in sets of vectors with similar relative positions among each other, i.e. similar cosine angles between node pairs, but they generally do not retain their absolute values (Dev et al., 2019; for the stability of the relations among embeddings see Wang et al., 2020). As a result, utilizing the CE framework to explore individual differences among subjects requires a method which would align different CEs to the same latent space (Fig. 1). The following sections evaluate our proposed embedding alignment method.

#### 3.1.1 Intra-individual CE similarity evaluation

We examined the effect of embedding alignment on the similarity of two independent CE fitting iterations of the same subject. This was done by testing the cosine similarity among pairs of identical nodes (Fig. 2a,i). Next, we examined embedding alignment effects on the similarity of the relations among pairs of different nodes, again, across two independent fittings of the same subject. The relations were quantified as the cosine similarity between their displacement vectors (Fig. 2b,i). **DS1**: For every subject (n = 100) the effect of embedding alignment on the mean cosine similarity of pairs of identical nodes (200 pairs) was tested. Significant improvement in node pair similarity (t(199) = 111.14, p<2.2e-16) was observed after applying embedding alignment (cos(θ)=0.98 ±0.001) compared to before alignment (cos(θ)=0.92 ±0.005). In contrast, the relations among pairs of different nodes, were highly consistent across independent fitting iterations both before (cos(θ)=0.99±2.7e-6) and after embedding alignment (cos(θ)=0.99±2.4e-5; Fig. 2b). Applying the same approach to **DS2** resulted in a similar outcome (t(232)= 118.31, p<2.2e-16; cos(θ)=0.96 ±0.004; cos(θ)=0.90 ±0.005; cos(θ)=0.99 ±8.7e-5; cos(θ)=0.99 ±9.1e-5; see SI 2 and SI Fig. 1a,b). These findings show that the alignment procedure successfully reduces the intra-individual variance in the embeddings. This is a crucial step towards identifying differences in CE embeddings obtained from different individuals.

**Fig. 2.**
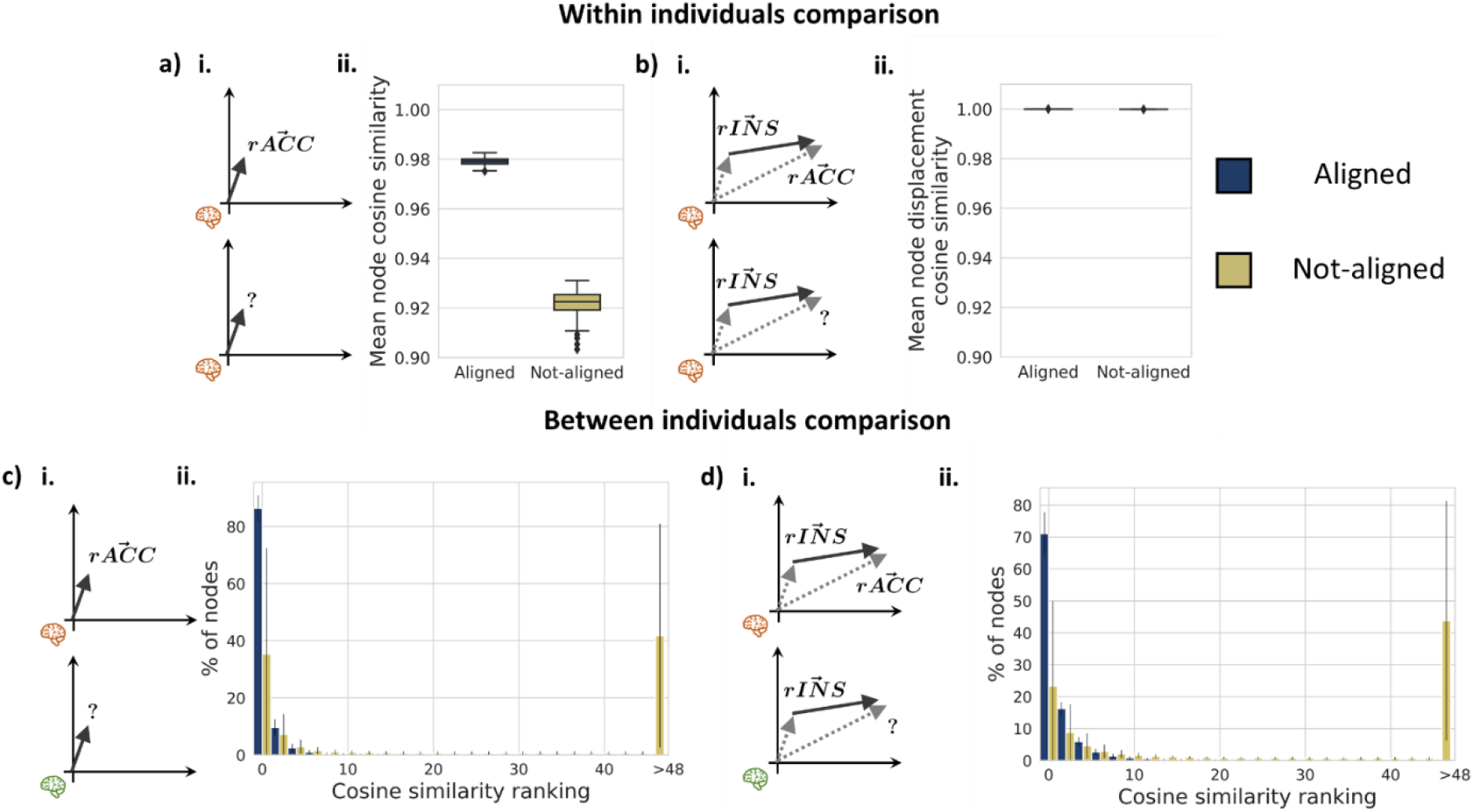
Individual CE alignment evaluation in DS1. Testing the effect of applying embedding alignment (dark blue) compared to its absence (yellow) on the similarity of identical nodes (a,c) and the similarity of displacement vectors among node pairs (b,d) across different embeddings. a, b Within individual comparisons. a.i.*Test 1* – the cosine similarity between identical nodes of the same individual across independent CE fitting iterations. a.ii. Box plot depicting the effect of embedding alignment on the distribution of the intra-individual node similarity for all subjects. b.i *Test 2* – the cosine similarity between the displacement vectors of all possible node pairs of the same individual across independent CE fitting iterations. b.ii. Box plot depicting the effect of embedding alignment on the distribution of the intra-individual nodes’ displacement vector similarity for all subjects. As is evident, across independent fitting iterations the cosine similarity among pairs of nodes is relatively stable while comparing individual nodes introduces considerable variation that could be reduced with CE alignment. c, d Between individuals comparisons. c.i.*Test 3* – the cosine similarity rank between identical nodes across different individuals. Cosine similarity rank was evaluated by ranking how close a given node in one individual is to the same node in a different individual. For example, a rank of 0 means that the closest neighbors of a given node, in terms of cosine similarity, across two individuals is itself. d.i. *Test 4* -the cosine similarity rank of an inter-individual nodes’ analogy test of all possible node pairs. For example, a rank of 0 means that the rACC of subject *a* minus the rINS of subject *a* plus the rINS of subject *b* was closest to the rACC of subject *b*. The effect of alignment on the ranking test was examined across all nodes (c.ii.) and all possible nodes pairs (d.ii.) Rank distributions are presented in bins of two; error bars represent the standard deviation across all subject pairs. The rINS and rACC were used only to illustrate the different tests conducted in the left panel of each plot, but the tests were conducted on all possible nodes and nodes pairs.

#### 3.1.2 Inter-individual CE similarity evaluation

Embedding alignment is meant to reduce variance that resulted from the stochasticity in node2vec fitting process. However, we do not expect perfect correspondence across individuals due to inherent differences in their underlying brain anatomy. To take this into account, we applied a more lenient test for estimating inter-individual CE similarity, the embedding ranking test (Rosenthal et al., 2018). Using the ranking test, we examined whether pairs of anatomically corresponding nodes across different individuals would be more similar than pairs of non-corresponding nodes, following embedding alignment. In this test, a given node would be ranked 0 if its closest neighbors in another individual, is the same anatomically corresponding node (Fig. 2c,i). In all analyses, query of the *i-*nearest node was done using cosine similarity. **DS1**: For every possible subject pair (n = 100; 9,950 pairs), we tested the effect of embedding alignment on the distribution of the node similarity rank. Before embedding alignment, 35.3% ±37.1% of the nodes were ranked in the top 2 nodes, compared to 86.3% ±4.6% following alignment. The percent of nodes ranked in the top 2 nodes was significantly higher following embedding alignment (t(9,949) = 135.7, p<2.2e-16; see Fig. 2c for the complete distribution). Similar results were found in **DS2** (n = 100; 9,950 pairs; 10.8% ±16.7%; 52.7% ±7.9%; t(9,949) = 246.6, p<2.2e-16; see SI 3 and SI Fig. 1c).

Next, to test the similarity of nodes’ relations we applied an analogy ranking test. This test measures whether the displacement vector from node *a* to *b* in one individual would express the same relation in another individual. We take the vectors of the right anterior cingulate cortex (rACC) and the right insula (rINS) as an example. In this case, a rank of 0 means that the rACC of subject *a* minus the rINS of subject *a* plus the rINS of subject *b* was closest to the rACC of subject *b*. **DS1**: The analogy test was conducted for every subject pair (a subset of n=30) and node pair (870 and 39,800 pairs respectively). Before embedding alignment 23.3% ±26.6% of the nodes were ranked in the top 2 nodes compared to 71.0% ±6.7% following alignment. This difference was significant (t(869) = 52.2, p<2.2e-16; Fig. 2d). Similarly in **DS2** (870 and 49,506 respectively; 4.1% ±6.5%; 31.2% ±5.2%; t(869) = 104.7, p<2.2e-16; see SI 3 and SI Fig. 1d).

### 3.2 Mapping structural to functional connectivity using CE

Resting-state functional connectivity refers to the temporal statistical dependence among activations in different brain nodes measured in the absence of an explicit task. While these patterns of coactivations depend on the structural connectivity, the observed correlation between the strengths of structural and functional connections is moderate, capturing only a fraction of the observed variance (Honey et al., 2010; Suárez et al., 2020). Testing this relation on the individual level, rather than the group level, further weaken the observed structure-function correspondence (Straathof et al., 2019). The CE framework was shown to improve the mapping of structural to functional connectivity at the group-level, presumably due to its ability to capture high-order graph relations (Rosenthal et al., 2018). Here we first reproduce those findings and extend the approach to structure-function relations within individuals.

#### 3.2.1 Group-level structure to function mapping

The relation between the group-level structural connectivity and the functional connectivity was quantified using Spearman’s rank correlation among corresponding functional and structural edges. The structural consensus connectivity matrix is typically sparse and accordingly, 38.9% and 18.9% of the group-level structural edges in DS1 and DS2, respectively, were larger than zero. For this reason, the correlation was taken only for node pairs linked by a structural edge, i.e. direct edges, in comparison to edges not linked by a structural connection, i.e. indirect. An additional measure of structural relation among nodes could be derived by taking the cosine angle among each pair of node embeddings. The CE cosine similarity was previously shown to substantially improve structure-function mapping (Rosenthal et al., 2018) and allowed to estimate this relation for both direct and indirect edges, as the cosine angle could be quantified even for pairs of nodes for which no connecting tracts were reconstructed. **DS1**: A significant correlation between the raw, direct structural edges and their corresponding functional connectivity edges was found (ρ(7782)=.311, p<2.2e-16; Fig. 4a). This structure-function correlation was higher when using the CE cosine similarity instead of the structural edges (ρ(7782, 12114, 19898)=.345,.137,.282; all p’s p<2.2e-16; for the direct, indirect and all edges respectively; Fig. 4b). Similar results were found in **DS2** (ρ(5032)= 0.287, p<2.2e-16; ρ(5032, 21992, 27026)=.389,.256,.340; all p’s p<2.2e-16; see SI 4 and SI Fig. 4a,b).

Next, we explored the contribution of individual edges to the CE-based structure-function mapping. Such estimation could be done on each unique edge since the CE cosine matrix is dense compared to the sparse raw structural connectivity matrix. We computed the structure-function correlation before minus after the removal of individual nodes. The result, the Δ correlation, informs us about the sign and relative magnitude of the contribution of each edge to the overall correlation (Gluth & Meiran, 2019; Fig. 3a, SI Fig. 2a). To test whether the Δ correlation could be solely attributed to the functional or the CE cosine edge values, we reported its correlation to the latter two. We compared the Δ correlation for direct versus indirect edges and for edges within the 7 canonical resting-states networks (Yeo et al., 2011) compared to between (see SI 4 for each of the networks). We additionally tested Δ correlation relation to Euclidian distance among node. **DS1**: A significant, weak correlation was found between the Δ correlation and the CE cosine (ρ(19898)=0.242, p p<2.2e-16) and the functional connectivity (ρ(19898)=0.167, p<2.2e-16) values. The Δ correlation was significantly larger for direct compared to indirect edges (t(19898)= 20.044, p<2.2e-16), and for edges within, compared to between canonical networks (t(19898)= 27.818, p<2.2e-16). Additionally, we found a significant interaction between the two factors (F(3,19896)= 1018.535, p<2.2e-16) such that, within compared to between networks’ edges were larger for direct edges (t(19898)= 39.943, p<2.2e-16) and a smaller, opposite effect was found for indirect edges (t(19898)= −16.586, p=1.0e-08; Fig. 3c). A small correlation was found between the Euclidian distance and Δ correlation (r(19898)=0.057, p<1.2e-15). Similar results were found in **DS2** (r(27026)=0.107, p=<2.2e-16, r(27026)=0.097, p<2.2e-16; t(27026)=26.392, p<2.2e-16; t(27026)=16.793, p<2.2e-16; F(3,27024)=762.826, p<2.2e-16; t(27026)=28.650, p<2.2e-16; t(27026)=-8.132, p =4.4e-16; r(27026)=0.115, p<2.2e-16; see SI 4, SI Fig. 2c and SI Fig. 3b). These results suggest that structure-function correspondence is largely driven by direct, within network edges.

**Fig. 3.**
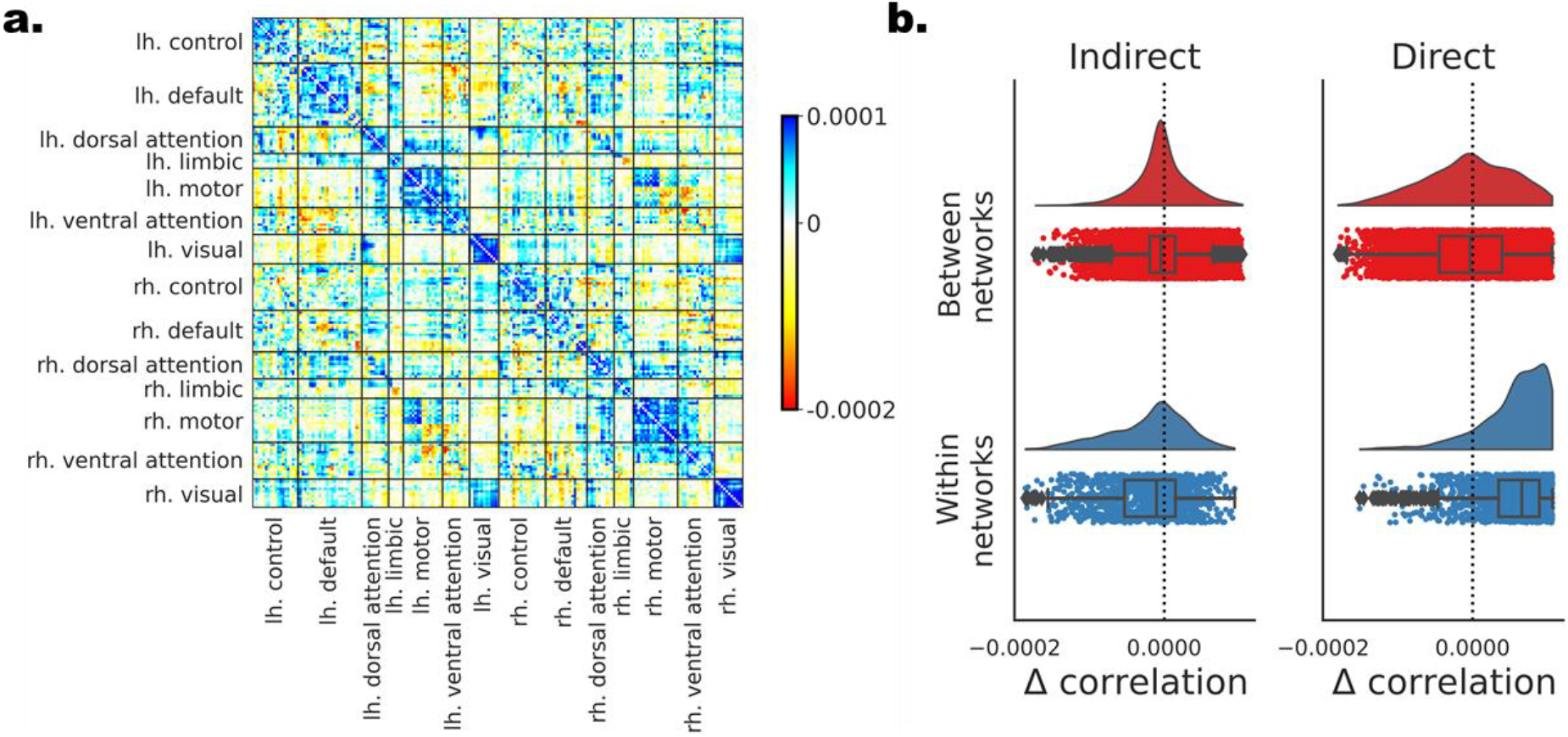
The contribution of individual edges to the CE-based structure-function mapping in DS1. Individual edges are assessed based on the difference in the measured structure-function correlation after removing each of the edges. (a) All edges are depicted in a matrix form. Nodes are ordered according to their affiliation to the 7 canonical resting-state networks (Yeo et al., 2011) separately for the left and right hemispheres. Edges with positive contribution to the overall correlation (blue) are those whose elimination results in lower correlation, while edges with negative contribution (red) have the opposite effect. (c) The distribution of Δ correlation for the edges is depicted for all 4 combinations of direct versus indirect and within versus between the 7 networks.

#### 3.2.2 Predicting group-level functional from structural connectivity with deep learning

The improvement in structure to function mapping using the CE cosine similarity was obtained without attempting to optimize this measure to match functional connectivity. In previous work (Rosenthal et al., 2018), this mapping was further improved using a deep learning model in which CEs were utilized as features to predict the functional edges. Specifically, the element-wise multiplication of pairs of nodes embeddings was used as input, and the predicted label was the observed functional connectivity among the two. We implemented a 4-layer, fully connected neural network, and the prediction was evaluated using cross-validation (see Method section). DS1: The Spearman’s rank correlation coefficient between the predicted and observed functional connectivity values were ρ(7782)=.641, ρ(12114)=.527 and ρ(19898)=.599, for the direct, indirect and all edges respectively, all p’s p<2.2e-16 (see Fig. 4c). Similar results were found in DS2 (ρ(5032)=.542, ρ(21992)=.488 and ρ(27026)=.527; all p’s p<2.2e-16; see SI 5 and SI Fig. 4c).

**Fig. 4.**
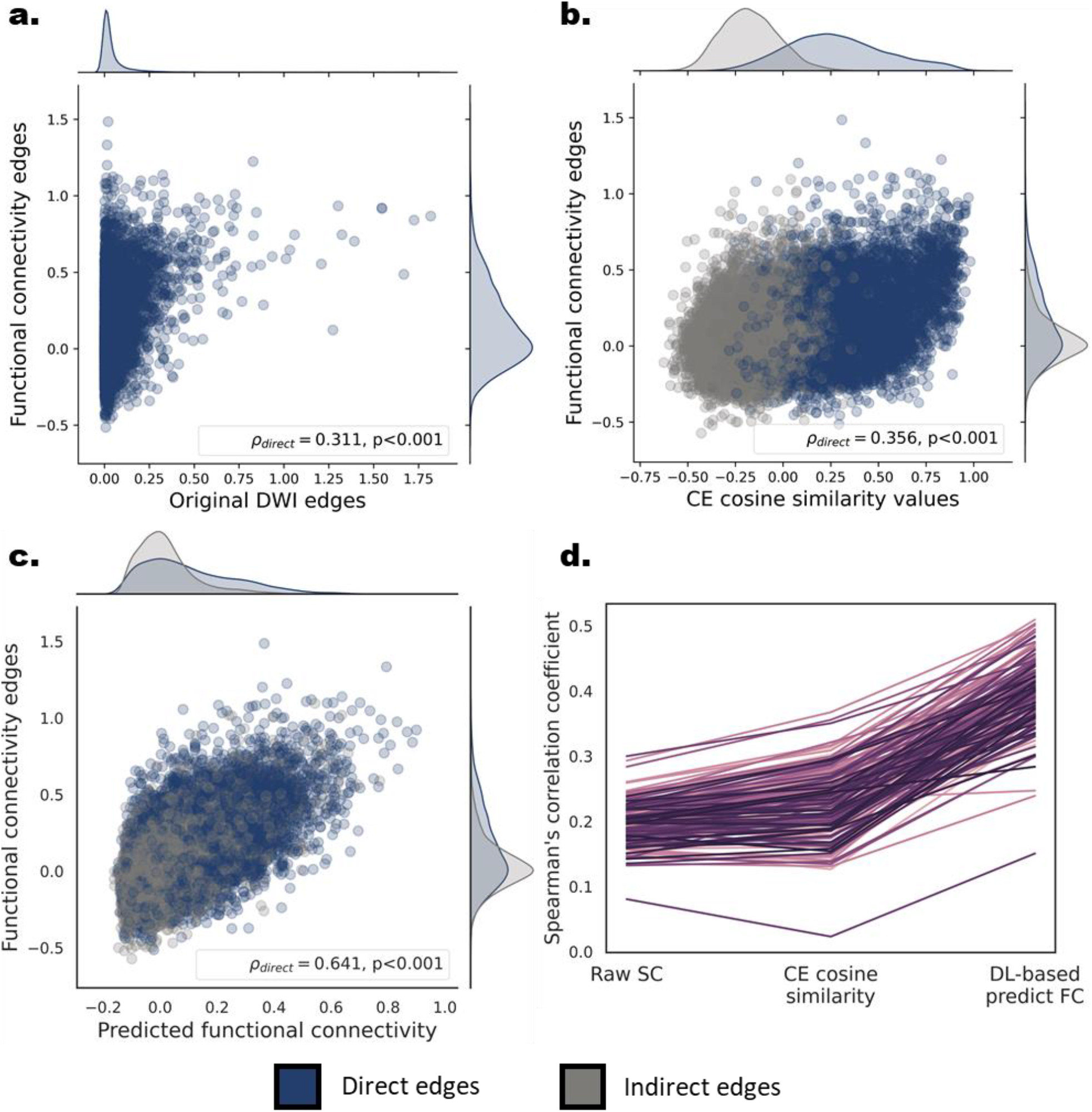
Correspondence between structural-functional connectivity at the group and individual levels in DS1. (a,b,c) Scatter plots and marginal univariate distributions of functional compared to structural-based edges at the group level. The structure-based edges are the entries of the streamline density matrix (a), the CE cosine similarities (b) and the deep learning based predicted functional edges. Direct edges are presented in dark blue and indirect edges in gray. The Spearman’s rank correlations and the corresponding p values for the group-level direct edges are depicted in the bottom right corner of each scatter plot. The correlations for the indirect edges were.137,.51 for the CE cosine similarities and the deep learning based predicted functional edges respectively. (d) Individual-level structure-function Spearman’s correlation values for direct edges of all three structure-based connectivity measures. Each line represents the correlation values of a single subject (n = 181). A significant increase in correlation was found for the CE cosine similarities compared to the structural edges and for the CE-based predicted functional edges compare to the CE cosine similarities.

#### 3.2.3 Individual-level structure to function mapping

Next, we tested whether the CE framework could be similarly utilized to map structural to functional connectivity within individuals. Structure-function correspondence was evaluated using Spearman’s rank correlation in each subject. We report this correlation using the structural connectivity values for direct edges and then again using the CE cosine similarity measure for direct and indirect edges. **DS1**: A significant correlation between the direct structural edges and their corresponding functional connectivity edges was found for all subjects (ρ=.197 ±.031, all p’s<2.2e-16; Fig. 3d). Similar matching to the CE cosine similarities values of direct edges revealed a significantly higher correlation (t(180) = 16.7, p<2.2e-16). The mean and standard deviation across subjects was ρ=.228 ±.048, ρ=.066 ±.035, ρ=.185 ±.037 for the direct, indirect and all edges, respectively. We found similar results in **DS2** (ρ=.152 ±.028, all p’s<2.2e-16; t(144) = 34.4, p<2.2e-16; ρ=.245 ±.040, ρ=.102 ±.035, ρ=.181 ±.344; see SI 6 and SI Fig.4d)

Finally, as in the group-level, we trained a deep learning neural network for predicting functional connectivity values from aligned CE, this time within individuals. **DS1**: A significant increase in the observed correlation for the direct edges was evident both compared to the structural edges (t(180) = 66.0, p<2.2e-16) and the CE cosine similarity measures (t(180) = 53.4, p<2.2e-16; SI Fig. 2d). The mean and standard deviation of the predicted, compared to the observed FC correlation was ρ=.397 ±.051, ρ=.253 ±.059, ρ=.337 ±.051, all p’s<2.2e-16, for the direct, indirect and all edges respectively. The same results were also obtained in **DS2** (t(144) = 52.8, p<2.2e-16; t(144) = 34.5, p<2.2e-16; ρ=.312 ±.043, ρ=.210 ±.051, ρ=.265 ±.046, all p’s<2.2e-16; see SI 6 and SI Fig. 4d)

#### 3.2.4 Parameters of the random walk and their relation to structure to function

The use of random walks to learn embeddings of brain nodes is strongly related to diffusive models of communication in the brain (Avena-Koenigsberger et al., 2018). Here we examine how such a diffusive process, as captured by the node2vec embedding algorithm, can model the relation between structural connectivity and functional connectivity. We adjusted the parameters of the random walk to be more globally or locally biased and measured the observed correlation between the CE cosine matrix and the functional connectivity. The parameters of the random walk were shifted from locally biased (p = 10^−3^, q =4.096) through unbiased (p = 1, q =1) to globally biased (p = 10^3^, q =0.244) random walks in 20 equal-spaced steps on a logarithmic scale. All results are present for the direct, indirect and all edges, respectively. **DS1**: Shifting from most locally biased (ρ=0.128±0.128, ρ=0.007±0.007, ρ=0.061±0.061) to the unbiased random walk (ρ=0.253±0.253, 0.082±0.082, 0.200±0.200) we observed a significant increase in structure to function mapping (t(11.5)=−17.033, −7.987, −18.184; all p’s<3.25e−08). Moving to the most globally biased random walk (ρ= 0.245±0.245, 0.068±0.068, 0.192±0.192) revealed a smaller but significant decrease in structure to function mapping (t(11.5)= −4.651, −5.564, −6.137; all p’s<1.01e−04; Fig. 5). In **DS2,** we found a similar increase when shifting from the most local to an unbiased random walk, but no significant difference moving to the most globally biased random walk (ρ= 0.202±0.202, 0.002±0.002, 0.049±0.049; ρ= 0.267±0.267, 0.122±0.122, 0.202±0.202; t(11.5)= −12.207, −20.914, −28.666; all p’s<8.76e−12; ρ= 0.262±0.262, 0.119±0.119, 0.200±0.200; t(11.5)= −2.789, −1.402, −1.267; p=0.010, 0.174, 0.217; see SI 7 and SI Fig. 5).

**Fig. 5.**
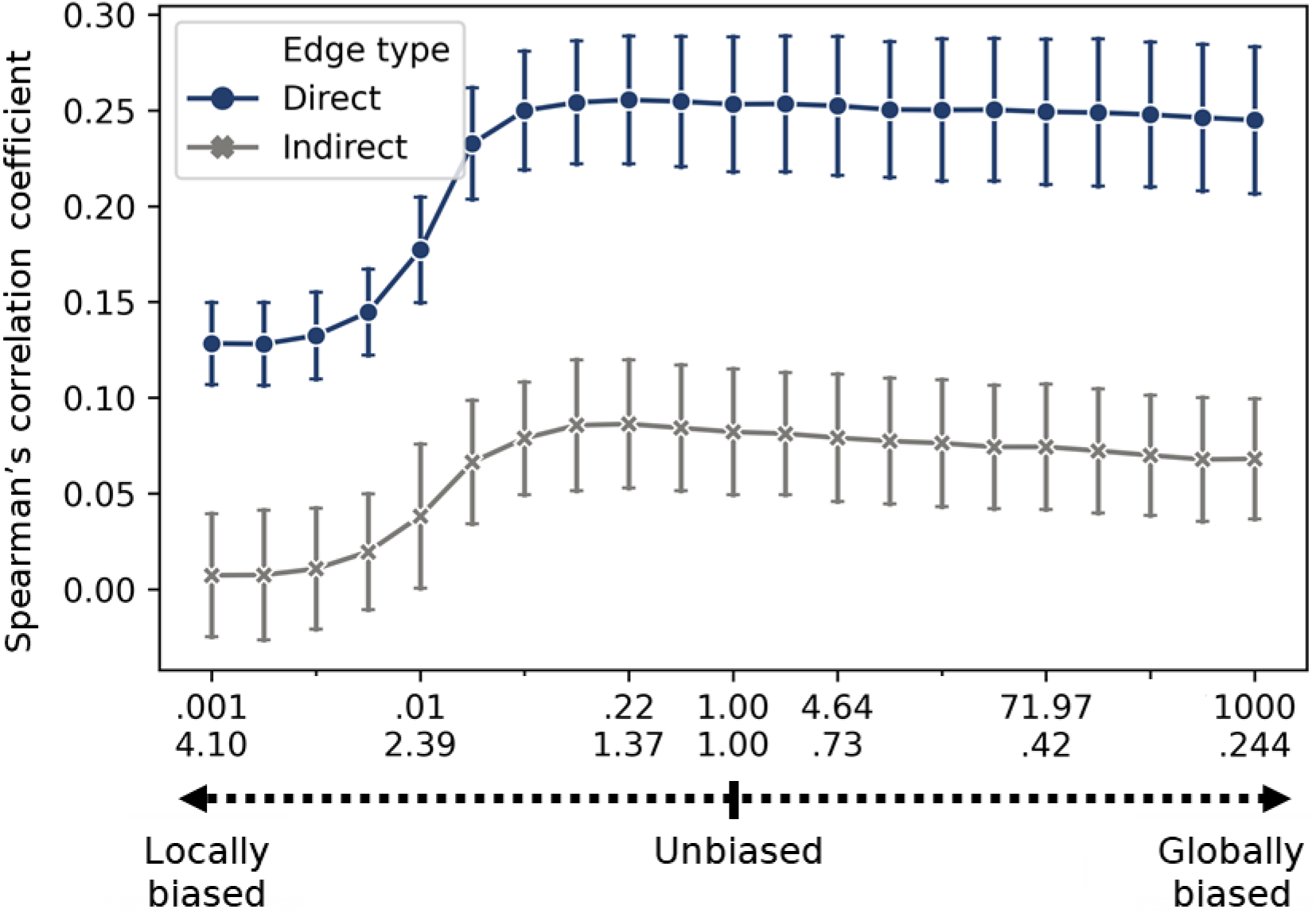
Effect of random walk parameters on structure-function correspondence in DS1. Testing the effect of random walks parameters on the Spearman’s rank correlation between direct (blue) or indirect edges (gray) to their corresponding functional connectivity edges. The random walk parameters were shifted from local (p = 10^−3^, q =4.096) through an unbiased (p = 1, q =1) to global (p = 10^3^, q =0.244) random walk in 20 equal bins on a logarithmic scale. Error bar represents the standard deviation across subjects (N = 25).

#### 3.2.5 Age-related changes in individual-level structure-function correspondence

Previous work reported age-related changes in structure-function coupling, but these were quantified at the whole connectome (Betzel et al., 2014) or the nodal (Zimmermann et al., 2016) levels. Here we examine whether the CE framework could be utilized to explore these alterations by testing the relation of aging to structure-function correlation. Importantly, the analysis was conducted both at the network level and at the edge-level by applying our edgewise contribution score (section 3.2.1) for each subject. **DS1**: A significant Pearson correlation was found between subjects’ age and structure-function correlation when using the structural edges (r(179)=−.406, p=1.4e−8) and the CE cosine similarity r(179)=−.487, p=3.6e−12; Fig. 6a). The observed increase in the correlation for the CE cosine similarity was significant (t=2.621, p=0.009). Next, we aimed to identify edges whose contribution to the structure-function correlation increases or decreases with age. We computed the edgewise contribution score for all subjects and correlated each edge with age (Fig. 6b). Using network contingency analysis, we revealed significant widespread age-related alteration in the structure-function contribution score (Fig. 6c). Examining edges within the 7 canonical functional networks (Yeo et al., 2011), we found a general decrease with age within hemispheres (t’s=6.94, 5.58, 3.02, 1.66; p’s = 5.9e−12, 2.82e−08, 0.002, 0.097; for |r| threshold of .25, .3, .35, .4 respectively) and increase between hemispheres (t’s=6.34, 6.04, 3.54, 1.73; p’s = 3.03e−10, 1.87e−09, 4.14e−04, 0.084; for |r| threshold of .25, .3, .35, .4). A mixed pattern was found between functional networks. Similar results were obtained in **DS2**: (r(144)=-.240, p=.003; r(144)=-.533, p=5.3e−12; t=4.9, p=1.4e−5; t’s= 3.83, 2.48, 2.09, 1.19; p’s = 1.3e−4, 0.013, 0.037, 0.233; t’s=14.7, 12.9, 11.1, 9.04; all p’s<2.2e−16; see SI 8, SI Fig. 6 and SI Fig. 8).

**Fig. 6.**
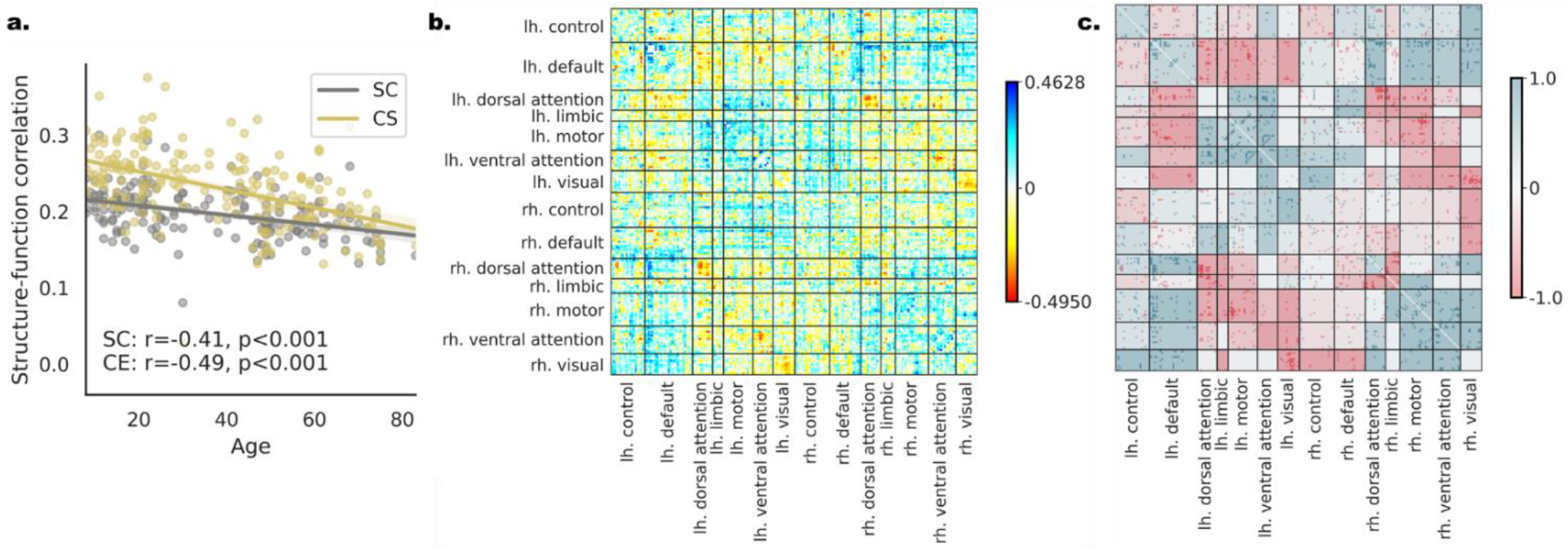
Age-related changes in individual-level structure-function correspondence in DS1. (a) A regression plot for age and the measured connectome-level structure-function correlation. The values of each subject are presented both for the structural edges (SC, gray) and the CE cosine edges (CE, yellow). The observed correlation to age is indicated on the bottom. The difference in the structure-function correspondence to age is significantly larger for the CE cosine connectivity measure. (b) The correlation of age to the edgewise contribution score of structure-function correspondence depicted in matrix form. Positive edges are ones that increase the overall correlation with age (blue) and negative edges are ones that decrease the overall correlation with age (red). (c) The result of the network contingency analysis for |r| > 0.25 (see SI Fig. 7 for all thresholds) in matrix form. Colored cells had a larger number of edges correlated with age than expected by chance. The cells’ background color is determined by the ratio between edges that are positively (blue) or negatively (red) correlated with age.

### 3.3 Modeling individual differences in age and intelligence using CE

Producing individual-level CE requires learning a compact vectorized representation of brain nodes and aligning the representations obtained from different individuals to the same latent space. We wanted to examine whether these transformations preserve variance associated with individual differences and whether representing connectomes with CE improves our ability to estimate those individual differences. Specifically, we used CE for out-of-sample prediction of age and intelligence in two separate datasets. In the previous section we focused on age-related individual differences in structural-functional correspondence. Here, age was directly predicted from CE.

#### 3.3.1 Capturing individual differences with CE

To predict age and intelligence we used a linear regression with CE and the CE cosine matrix as input. First, within the training set, we used 5-fold CV to predict each of the two outcomes using each node as an input. Then, individual nodes’ predictions were combined both by fitting a second-level model or by taking their mean, and testing on the left-out test set. The performance of the models was quantified as the correlation between the observed and predicted age or intelligence. **DS1**: For age, the mean observed-predicted correlation across all nodes was r=.465(±.078) for CE and r=0.467(±0.057) for the CE cosine matrix. Similarly, for intelligence r=.093(±.070) for CE, and r=0.083(±0.066) for the CE cosine matrix. The connectome-level model, validated on the left-out test set, revealed a significant correlation of the observed-predicted age (CE: r(188)=0.770, 0.760, cosine CE: r(188)=0.826, 0.750; for the second-level model and the mean respectively; all p’s<2.2e−16) and intelligence (CE: r(188)=0.379, 0.265, cosine CE: r(188)=0.337, 0.200; for the second-level model and the mean respectively, all p’s<0.008; see Fig.7). Applying the same analysis to **DS2** yielded similar results (r=.453(±.078), r=0.367(±0.056); r=.240(±.077), r=0.138(±0.060); r(199)=0.801, 0.753, r(199)=0.836, 0.777; all p’s<2.2e−16; r(199)=0.578, 0.360, r(199)=0.565, 0.409; all p’s<3.5e−07; see SI 9.1 and SI Fig. 9). In both datasets all results were reproduced for age prediction after controlling for gender. Controlling for age and gender in intelligence prediction reproduced all results in DS1, while the observed effects were reduced in DS2, such that a small, yet significant effect was found for the ensemble model only for the CE cosine matrix as input (see SI 9.3).

**Fig. 7.**
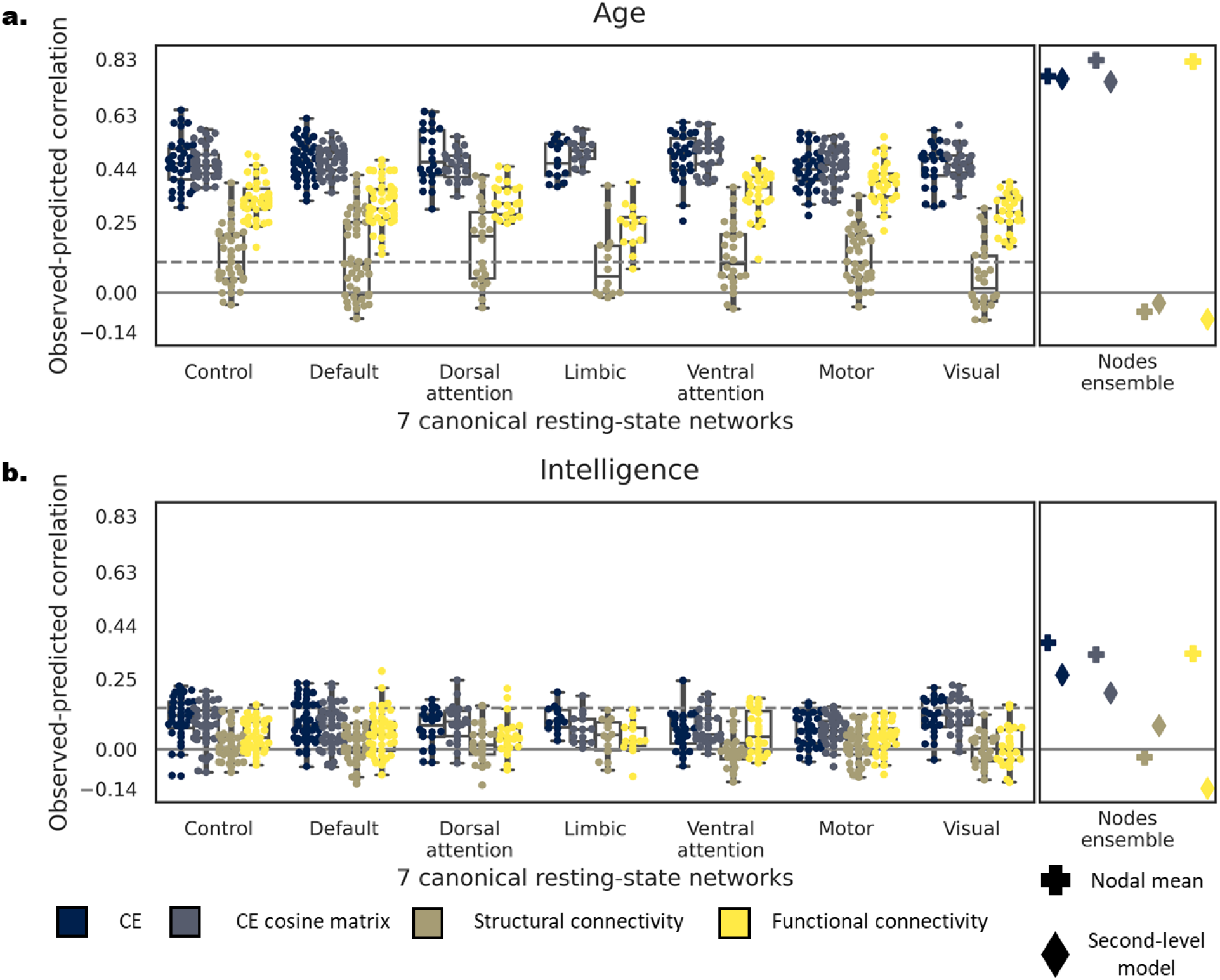
Comparing predictive accuracy of CE, CE cosine similarity matrix, structural and functional connectivity as input within DS1. Correlation between the observed and predicted age (a) and intelligence (b) with CE (blue), CE cosine matrix (blue-gray), structural (brown) and functional (yellow) connectivity. On the left panel of each subplot, dots represent the predictive accuracy of each node within each of the 7 resting state networks (Yeo et al., 2011). Nodes are grouped by their resting state network affiliation. The right panel depicts the predictive accuracy of a model combining all individual nodes by taking their mean (plus symbol) or aggregates them using a second-level linear model (diamond symbol). The dashed line represents the FDR-corrected significance level for single nodal prediction.

#### 3.3.2 Predictive accuracy of CE compared to structural and functional connectivity

Next, we examined whether predictive accuracy gained with CE would be superior compared to using only structural or functional connectivity as input. We repeated the same steps for the two additional inputs and compared the resulting performance to CE across all nodes using a paired t-test. **DS1**: Using both CE and the CE cosine matrix resulted in significantly better performance than functional and structural connectivity for age (all t’s(199) > 17.1, p’s <2.2e−16) and intelligence (all t’s (199) > 6.91, all p’s <6.6e−11). The structural connectivity connectome-level model, validated on the left-out test set, resulted in nonsignificant correlation for both age and intelligence (all r’s(188) < 0.084, p’s >0.267). Contrarily, functional connectivity resulted in significant correlation when taking the mean of the nodal predictions (age: r(188)=0.822, p’s <2.2e−16; intelligence: r(188)=0.341, p=3.7e−06) but not for the second-level model (all r’s(188) < 0.000; see Fig.7). Applying the same analysis to **DS2** yielded similar results (all t’s(232) > 4.36, all p’s <2.0e−05; all t’s (232) > 2.53, all p’s <0.012, except for cosine embedding and functional connectivity (t(232) =−0.947p: 0.345); all r’s(137) < 0.005, p’s >0.415; r(137)=0.854, p’s <2.2e−16; r(137)=0.546, p=4.4e−12; all r’s(137) < 0.039; see SI 9.2 and SI Fig. 9). In both datasets all results were reproduced for age prediction after controlling for gender. Controlling for age and gender for intelligence prediction reproduced all results in DS1 but not in DS2 (see SI 9.4). This might result from a smaller sample size of individuals who had both structure and functional data in DS2 (425 compared to 601 subjects).

## 4. Discussion

Studies of the human connectome have added to our understanding of the organizing principles of neural processing and communication in the brain. However, the relation between observed individual differences in connectome topology and individual differences in function and behavior is still poorly understood. In the current work, we leverage the CE framework for modeling individual differences by aligning embeddings of different individuals to a common space. We empirically evaluated our alignment scheme and found a large increase in node’s similarity across subjects, indicating successful alignment of individual connectome embeddings. We then demonstrated two contributions of applying the CE framework at the individual subject level. The first is the improvement in subject-level mapping of structural to functional connectivity while preserving age-related variance associated with this structure-function correspondence. The second contribution is the successful use of aligned CE in predicting demographic and behavioral variables. CE resulted in significantly improved prediction of age and intelligence compared to structural or functional connectivity alone, suggesting that CE not only preserves variability related to demographics and behavior, but also accentuates this variability such that it is more accessible for prediction-based models.

### 4.1 CE alignment

Enabling the mutual alignment of CEs is a necessary step to utilize machine learning techniques for subsequent tasks such as prediction of functional connectivity or the presence of a neurological condition. Here we consider two problems that may be aided by proper CE alignment. The first is to find an optimal one-to-one mapping between nodes taken from one connectome to another based on their topological attributes. Such mapping could be used, for example, to find homologous brain structures among different species. The second involves aligning vectorized representations of connectomes in the same latent space to allow their comparison. Here, the mapping of corresponding nodes across connectomes is known *a priori* and we are interested in the subtle variations in these vectorized representations that arise due to differences in the sampled random walks. The goal is to remove variability related to the stochasticity in the CE fitting process and the different latent spaces, and preserve variability related to variations in the underlining network topology. While the first problem has been previously addressed in applications such as translation among languages (Smith et al., 2017), to the best of our knowledge, the second has not been addressed before. We suggest that our CE alignment method could additionally be used in linguistic tasks such as authorship attribution (Kocher & Savoy, 2018) or examining differences in word semantics among different cultures (Karimi et al., 2015). Notably, our method is computationally efficient as it does not require the co-learning of all embeddings when encountering a new sample (Wolf et al., 2014) making it more desirable for clinical applications.

### 4.2 Mapping Structural to functional connectivity

The observed correlation between group-level structural and functional connectivity is typically moderate (r = 0.3-0.5; Suárez et al., 2020). In part, moderate correlations may reflect limits on acquisition and reconstruction, as well as the fact that the same structural backbone supports a large and dynamic repertoire of functional interactions (Deco et al., 2013; Fukushima et al., 2018). Nevertheless, considering high-level structural interactions among nodes might explain a larger portion of the functional connectivity variance. In the current work, we demonstrate that using a deep learning model, functional connectivity could be predicted from CE, while substantially improving structure-function mapping at the group-level (DS1: ρ=0.64, DS2: ρ=0.54). The observed structure-function correlation at the individual subject level is often more modest than group-level estimates (r = 0.02-0.25; Straathof et al., 2019) possibly due to noise associated with the acquisition of the different imaging modalities, under-sampling of resting-state dynamics in short scans (Birn et al., 2013), and variation in functional boundaries among individuals (Gordon et al., 2017). Applying the CE approach to similar mapping at the individual-level resulted in significant improvements of structure-function correlations (DS1: ρ=0.397, DS2: ρ=0.312). Additionally, CE produced an edge-wise similarity measure even for pairs of nodes for which no connecting tracts were found. This allows a more comprehensive estimate of edge-wise contribution to structure-function correspondence. Our results suggest that a specific subset of edges (direct edges within canonical resting-state networks) drive the observed structure-function correlation. Finally, we showed that CE preserved and even enhanced age-related individual differences in structure-function mapping.

### 4.3 CE as a model of communication linking structure to function

CE may advance our understanding of the nature of communication in the brain. Communication or information flow in brain networks could be seen as the complete set of dynamic causal influences among pairs of neuronal elements (Avena-Koenigsberger et al., 2018). According to this view, communication is constrained by the structural connectivity scaffold and gives rise to the observed co-fluctuations among pairs of brain regions (functional connectivity), and hence it offers a link between the two. Models of communication processes among brain regions range from diffusive flow of information, as in random walks, to routing mechanisms, such as schemes based on shortest paths. CE, fitted based on a set of random walks, can be viewed as a generative model of diffusive communication around nodes. Their success implies that diffusive models might better capture the observed functional connectivity (Goni et al., 2014). Further manipulation of the specific random walk parameters used to fit the CE model, suggest that an unbiased, or a slightly locally biased random walk best accounts for the observed functional connectivity. In future work the sampled walks could be generated based on different mechanisms (e.g. shortest paths) and their relation of corresponding embeddings to static or dynamic connectivity could be tested.

### 4.4 Prediction of individual differences using aligned CE

Cross-subject alignment, whether conducted based on anatomy (Frost & Goebel, 2012) or function (Haxby et al., 2020), is a critical step for highlighting individual differences in brain mapping. Here we consider the characterization of individual differences as an ideal test case for the CE alignment approach. Indeed, we found that aligned CE, as well as CE cosine similarity matrices show a robust increase in prediction accuracy compared to structural connectivity for age and intelligence both at the individual node and the whole connectome levels. This observed increase in predictive accuracy could be attributed to the sparsity and dimensionality of the structural connectivity matrix, and its inability to capture high-order topological relations. These might explain how despite a growing trend in studies focusing on individual differences (Sui et al., 2020), fewer studies so far have applied a predictive modelling on structural connectivity data as compared to other modalities, such as functional connectivity (Arbabshirani et al., 2017). Our CE framework addresses these issues by learning node representations that are low dimensional and preserves nodes’ topological context, rather than its mere direct connections. When depicted in the form of a cosine similarity matrix it captures all pair-wise relations among nodes resulting in a dense matrix representation. In addition to this advantage in representing structural connectivity, CE can be viewed as a model of brain communication. Thus, it represents a functional aspect of brain connectivity in addition to structure. This might explain the advantage at nodal-level or the similarity for the connectome-level model of CE compared to functional connectivity in predictive accuracy. Note that functional connectivity exhibits comparable results only for one of the connectome level models (nodal mean). Thus, the CE framework might hold promise to advance the application of connectome mapping to such predictive models. Future work can utilize the CE predictive framework to study connectome alterations in neurodevelopmental conditions. Furthermore, we can explore the effect of “lesioning” of the network by for example, zeroing out connections or entire nodes before the embedding process. Then, the effect of these “lesions” on the predicted outcome in such prediction models could be tested (Rosenthal et al., 2018)

### 4.5 Conclusions

Our findings suggest that learned connectome representations and their mutual alignment are a powerful tool for conducting individual-level mapping of structural connectivity to function and behavior. We suggest two complementary views on CE. The first, is an improved structural connectivity representation, allowing to quantify structural relations that are missing from the network reconstruction. The second, as a proxy for diffusive communication on the structural backbone, hence incorporating aspects of structure (the structural graph) as well as function (the diffusive process). The advancements made here in the CE framework support current efforts in neuroscience to better understand and address individual differences.

## 5. Data availability

The unprocessed data is openly available online at http://fcon_1000.projects.nitrc.org/indi/enhanced/neurodata.html for DS1and available upon online access request https://camcan-archive.mrc-cbu.cam.ac.uk/dataaccess/ for DS2. A python package implementation of CE framework (https://github.com/GidLev/cepy) and set of interactive notebooks reproducing the main results of the paper (https://github.com/GidLev/cepy/tree/master/examples) were made available online.

## 6. Acknowledgments

This research was supported by the U.S.-Israel Binational Science Foundation (BSF), grant 2017242 to GA and OS. OS was partially supported by NIH grant R01MH122957. This material is based upon work supported by the National Science Foundation Graduate Research Fellowship under Grant No. 1342962 (J.F.).

