## Supplementary information for "Mapping individual differences across brain network structure to function and behavior with connectome embedding"

1. *MRI acquisition and preprocessing*

**Supplementary table 1. Acquisition parameters of the T1w, DWI, and rsfMRI for each dataset**

|  | NKI (DS1) | CamCan (DS2) |
| --- | --- | --- |
| Scanner | Siemens Magneton Trio | 3T Siemens Trio |
| Scanner strength | 3T | 3T |
| Head coil | 12-channel | 32-channel |
| T1w |  |  |
| TR | 1900 ms | 2250 ms |
| TE | 2.52 ms | 2.99 ms |
| TI | 900 ms | 900 ms |
| Flip angle | 9 | 9 |
| Field of view | 350 mm × 263 mm × 350 mm | 256 mm × 240 mm × 192 mm |
| Voxel size | 1 mm isotropic | 1 mm isotropic |
| DWI |  |  |
| TR | 2400 ms | 9100 ms |
| TE | 85 ms | 104 ms |
| Field of view | 212 mm × 180 mm × 128 mm | 192 mm × 192 mm × 142mm |
| B0 volumes | 9 | 3 |
| Gradient directions | 128 | 60 |
| b-values | 1500 s/mm^2^ | 1000, 2000 s/mm^2^ |
| Voxel size | 2 mm isotropic | 2mm isotropic |
| RsfMRI |  |  |
| TR | 645 ms | 1970 ms |
| TE | 30 ms | 30 ms |
| Flip angle | 60 | 78 |
| Field of view | 222 mm × 222 mm × 120 mm | 192 mm × 192 mm × 142 mm |
| Voxel size | 3 mm isotropic | 3 mm × 3 mm × 4.44 mm |
| Scan time | 9.68 minutes | 8.56 minutes |
| Instructions | Rest with fixation on cross | Rest with eyes closed |

1.1 *MRI preprocessing – DS1*

T1w scans were processed through FreeSurfer’s recon-all pipeline (version 6.0). The FreeSurfer cortical reconstruction function was used to skull strip the T1w, which was aligned to MNI space with 6 degrees of freedom. The Shaefer parcellations were transferred to T1w space using the spherical surface warp generated by FreeSurfer.

DWI were preprocessed following the DESIGNER pipeline (Ades-Aron et al., 2018), which includes MP-PCA denoising (Veraart et al., 2016), Gibbs ringing artifact correction (Kellner et al., 2016), Rician bias correction, distortion and eddy current correction (Andersson & Sotiropoulos, 2016), and B1 field correction (Veraart et al., 2016). DWI were then aligned to their corresponding T1w (7 degrees of freedom) and the MNI space in one interpolation step. B-vectors were rotated accordingly. Local models of white matter orientation were estimated in a recursive manner (Tax et al., 2014) by applying Constrained Spherical Deconvolution with a spherical harmonics order of 8. Probabilistic streamline tractography (Garyfallidis et al., 2014) was seeded five times in each white matter voxel and streamlines less than 10mm were discarded. Streamline counts between regions were normalized by the geometric mean volume of the two regions.

Resting-state preprocessing was performed using the fMRIPrep version 1.1.8 (Esteban et al., 2019). Parts of the following description have been reproduced from the fMRIPrep documentation. In this pipeline, each T1w was corrected using N4BiasFieldCorrection v2.1.0 (Tustison et al., 2010) and skull-stripped using antsBrainExtraction.sh v2.1.0 (using the OASIS template). The ANT-s derived brain mask was refined with a custom variation of the method to reconcile ANTs-derived and FreeSurfer-derived segmentations of the cortical gray-matter of Mindboggle (Klein et al., 2017). Brain tissue segmentation of cerebrospinal fluid (CSF), white-matter (WM) and gray-matter (GM) was performed on the brain-extracted T1w using fast (FSL v5.0.9; Zhang, Brady, & Smith, 2001). Functional data was slice time corrected using 3dTshift from AFNI v16.2.07 (Cox, 1996) and motion corrected using mcflirt (FSL v5.0.9; Jenkinson, Bannister, Brady, & Smith, 2002). "Fieldmap-less" distortion correction was performed by co-registering the functional image to the same-subject T1w with intensity inverted (Huntenburg, 2014; S. Wang et al., 2017) constrained with an average fieldmap template (Treiber et al., 2016), implemented with antsRegistration (ANTs). This was followed by co-registration to the corresponding T1w using boundary-based registration (Greve & Fischl, 2009) with 9 degrees of freedom, using bbregister (FreeSurfer version 6.0.1). Motion correcting transformations, field distortion correcting warp, and BOLD-to-T1w transformation warp were concatenated and applied in a single step using antsApplyTransforms (ANTs v2.1.0) using Lanczos interpolation. Frame-wise displacement (Power et al., 2014) was calculated for each functional run using the implementation of Nipype. The first four frames of the BOLD data in the T1w space were discarded. These data were then bandpass filtered (0.008 – 0.08Hz) and then confound regressed in a manner orthogonal to the temporal filters using 6 motion estimates, the mean time series derived in CSF, WM, and whole brain masks, the derivatives of these nine regressors, and the squares of these 18 terms (Parkes et al., 2018; Satterthwaite et al., 2013). Spike regressors were added for each frame with framewise displacement above 0.5mm. Data were linearly detrended and standardized.

1.2 *MRI preprocessing – DS2*

T1w were processed through FreeSurfer’s recon-all pipeline (version 6.0). The FreeSurfer reconstruction was used in tandem with the Connectome Mapper, to recover the Lausanne 223 node parcellation in T1w space.

DWI were corrected for eddy currents (Andersson & Sotiropoulos, 2016), denoised (Garyfallidis et al., 2014), motion-corrected (Jenkinson et al., 2002), and aligned to the corresponding T1w (Avants et al., 2012). Gray and white matter segmentation was based on FreeSurfer’s version 6 recon-all (Fischl et al., 1999, 2004). Constrained Spherical Deconvolution was used for deterministic streamline reconstruction (Garyfallidis et al., 2014).. The structural connectivity matrices indicated the number of streamlines connecting each pair of ROIs weighted by the mean volume of the two, i.e. connectivity density. Edges with less than 10 connecting streamlines were set to zero. Subjects for which more than 10 nodes in the structural connectivity matrix had a degree of 0 were excluded and were not used in subsequent analysis (6% omitted, 601 left). Resting-state preprocessing was performed using the Configurable Pipeline for the Analysis of Connectomes (C-PAC; Cameron et al., 2013) and included: slice time correction, motion correction, skull removal, regression of the first 5 principal components of signal from white matter and CSF (Behzadi et al., 2007), 6 motion parameters and linear and quadratic trends, global signal regression, followed by temporal filtering between 0.1 and 0.01 Hz and. Finally, a scrubbing threshold of 0.5mm frame-wise displacement (Power et al., 2014; removal of 1 TR before and 2 TR after excessive movement) was applied. Exclusion criterion for excessive movements was determined a priori to less than 50% (4 min and 20 sec) of the resting-state session after the scrubbing procedure (24% omitted; 490 subjects left).

Functional connectivity as used in all analyses was defined as the Pearson correlation among pairs of ROIs’ time series following Fisher’s r-to-z transformation. The samples of both datasets were divided into a training (67%) and test sets (33%) for subsequent tasks that required out-of-sample accuracy estimation (ds 1: 361 and 181, DS2: 401 and 200; training and test subjects, respectively).

1.3 *NKI exclusion criteria*

The NKI was downloaded in December of 2016 from the INDI S3 Bucket. At the time of download, the dataset consisted of 957 T1w (811 subjects), 914 dMRI (771 subjects), and 718 fMRI (“acquisition645”; 634 subjects) images. T1w and dMRI images, and tractography results were first filtered based on visual inspection. T1w images were filtered based on artifact, such as ringing or ghosting (43 images) and for FreeSurfer reconstruction failure (105 images), leaving 809 T1w images (699 subjects). dMRI images were filtered based on corrupt data (13 images) and artifact on fitted fractional anisotropy maps (18 images), leaving 883 images (747 subjects). Tractography was run on 781 images (677 subjects) that had both quality controlled T1w and dMRI images. Tractography results were filtered based on artifact, which include failure to resolve callosal, cingulum, and/or corticospinal streamlines or errors resulting in sparse streamline densities, leaving 764 tractography results (661 subjects). T1w, dMRI, and fMRI images were then filtered using computed image quality metrics (Esteban et al., 2017; Roalf et al., 2016). T1w images were excluded if the scan was marked as an outlier (1.5x the inter-quartile range in the adverse direction) in three or more of following quality metric distributions: coefficient of joint variation, contrast-to-noise ratio, signal-to-noise ratio, Dietrich’s SNR, FBER, and EFC. dMRI images were excluded if the percent of signal outliers, determined by eddy_qc (Bastiani et al., 2019), was greater than 15%. Furthermore, dMRI were excluded if the scan was marked as an outlier (1.5x the inter-quartile range in the adverse direction) in two or more of following quality metric distributions: temporal signal-to-noise ratio, mean voxel intensity outlier count, or max voxel intensity outlier count. fMRI images were excluded if greater than 15% of time frames exceeded 0.5mm framewise displacement. Furthermore, fMRI images were excluded the scan was marked as an outlier (1.5x the inter-quartile range in the adverse direction) in 3 or more of the following quality metric distributions: DVARS standard deviation, DVARS voxel-wise standard deviation, temporal signal-to-noise ratio, framewise displacement mean, AFNI’s outlier ratio, and AFNI’s quality index. This image quality metric filtering excluded zero T1w images, 16 dMRI images, and 21 fMRI images. Following visual and image quality metric filtering, 809 T1w images (699 subjects), 728 dMRI images (619 subjects), and 697 fMRI images (633 subjects). The intersection of subjects with at least one valid T1w, dMRI, and fMRI images totaled 567 subjects. Finally, age metadata was available for 542 of these subjects.

2. *Intra-individual CE similarity evaluation in DS2*

Significant improvement in the similarity of identical nodes (t(232) = 118.31, p<2.2e-16; Fig. S1.a) was observed after applying embedding alignment (cos(θ)=0.96 ±0.004) compared to its absence (cos(θ)=0.90 ±0.005). While the displacement vectors among all possible pairs of different nodes where highly consistent (cos(θ)=0.99 ±8.7e-5, cos(θ)=0.99 ±9.1e-5; with and without embedding alignment respectively; Fig. S1.b)

3. *Inter-individual CE similarity evaluation in DS2*

The effect of embedding alignment on nodes similarity rank distribution - For every possible subject pair (n = 100; 9,950 pairs), we tested the effect of embedding alignment on nodes similarity rank distribution. Before embedding alignment, 10.8% ±16.7% of the nodes were ranked in the top 2 nodes compared to 52.7% ±7.9% following alignment. Again, this difference was significant (t(9,949) = 246.6, p<2.2e-16; Fig. S1.c).

The analogy test - This test was conducted for every possible subject and node pairs (870 and 49,506 respectively). Before embedding alignment, 4.1% ±6.5% of the nodes were ranked in the top 2 nodes compared to 31.2% ±5.2% following alignment. The percent of nodes ranked in the top 2 nodes was significantly higher following embedding alignment (t(869) = 104.7, p<2.2e-16; Fig. S1d).

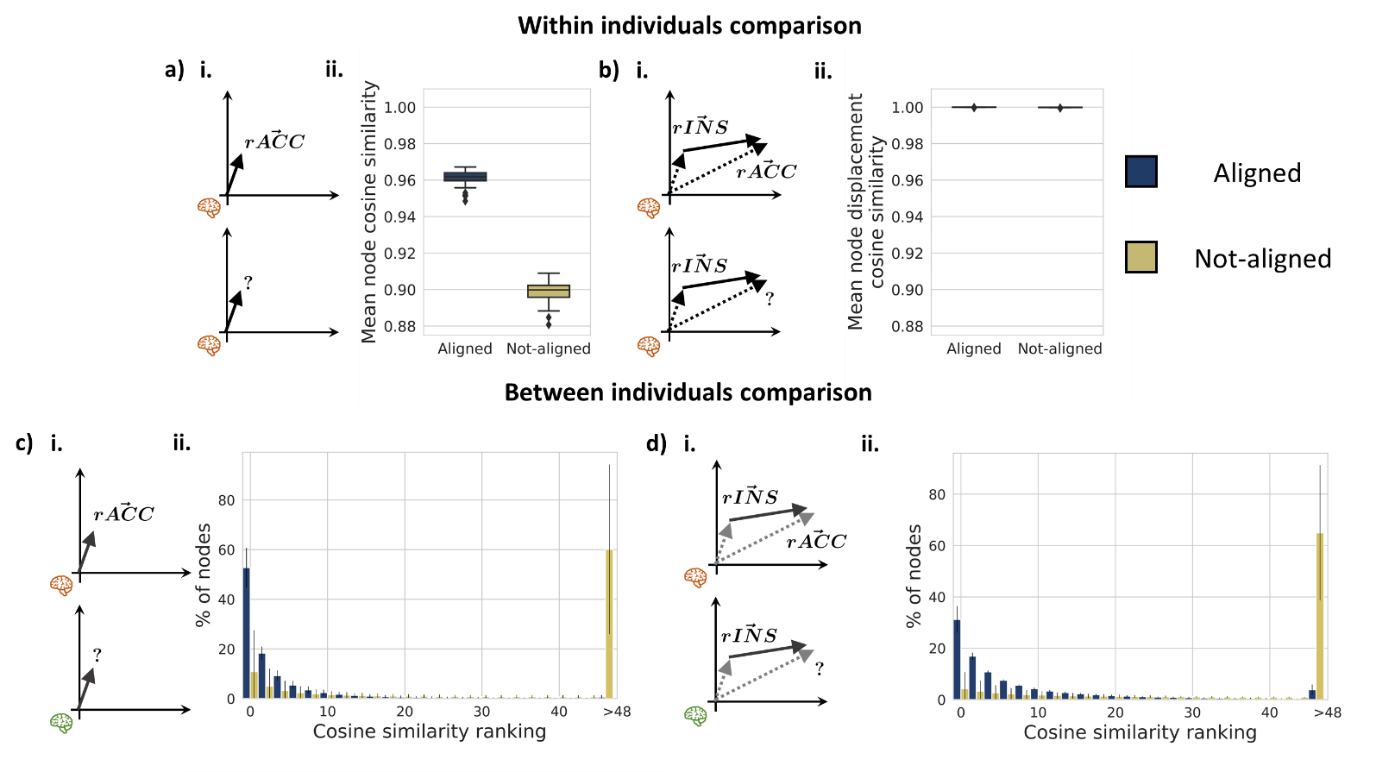

**SI Fig. 1. Individual CE alignment evaluation in DS2. Testing the effect of applying embedding alignment (dark blue) compared to its absence (yellow) on the similarity of identical nodes (a,c) and the similarity of displacement vectors among node pairs (b,d) across different embeddings. a, b Within individual comparisons a.i. *Test 1* – the cosine similarity between identical nodes of the same individual across independent CE fitting iterations. a.ii. Box plot depicting the effect of embedding alignment on the distribution of the intra-individual node similarity for all subjects. b.i *Test 2* – the cosine similarity between the displacement vectors of all possible node pairs of the same individual across independent CE fitting iterations. b.ii. Box plot depicting the effect of embedding alignment on the distribution of the intra-individual nodes’ displacement vector similarity for all subjects. As is evident, across independent fitting iterations, the cosine similarity among pairs of nodes is relatively stable while comparing individual nodes introduces considerable variation that could be reduced with CE alignment. c, d Between individuals comparisons c.i. *Test 3* – the cosine similarity rank between identical nodes across different individuals. Cosine similarity rank was evaluated by ranking how close a given node in one individual is to the same node in a different individual. For example, a rank of 0 means that the closest neighbors of a given node, in terms of cosine similarity, across two individuals is itself. d.i. *Test 4* - the cosine similarity rank of an inter-individual nodes’ analogy test of all possible node pairs. For example, a rank of 0 means that the rACC of subject *a* minus the rINS of subject *a* plus the rINS of subject *b* was closest to the rACC of subject *b*. The effect of alignment on the ranking test was examined across all nodes (c.ii.) and all possible nodes pairs (d.ii.) Rank distributions are presented in bins of two; error bars represent the standard deviation across all subject pairs. The rINS and rACC were used only to illustrate the different tests conducted in the left panel of each plot, but the tests were conducted on all possible nodes and nodes pairs.**

4. *Group-level structure to function mapping*
   4.1 *Edgewise contribution score across functional networks in DS1*

Examining each network separately we found a significant positive contribution to structure-function correlation for edges within the motor network (t(33)=2.535, p=0.016) and negative contribution for edges within the ventral attention network (t(25)=-2.252, p=0.033; Fig. 3b). Repeating the test only for edges within the functional networks reveled a significant positive contribution for edges within all networks (all t’s>2.472, all p’s<0.014; SI Fig 3a). A nodal average of Δ correlation is depicted on the brain surface (SI Fig. 3c,d).

4.2 *Connectome-level structure to function mapping in DS2*

Along with the results of DS1, a significant correlation between the direct structural edges and their corresponding functional connectivity edges was found (ρ(5032)= 0.287, p<2.2e-16; Fig. S2.a). The correlation was higher when comparing the CE cosine similarity instead of the structural edges (ρ(5032, 21992, 27026)=.389,.256,.340; all p’s p<2.2e-16; for the direct, indirect and all edges respectively; Fig. S2.b).

4.3 *Edgewise contribution score in DS2*

A significant, weak correlation was found between the Δ correlation and the CE cosine (r(27026)=0.107, p=<2.2e-16) and the functional connectivity (r(27026)=0.097, p<2.2e-16) values. The Δ correlation was significantly larger for direct compared to indirect edges (t(27026)=26.392, p<2.2e-16), and for edges within compared to between networks (t(27026)=16.793, p<2.2e-16). Additionally, we found a significant interaction between the two factors (F(3,27024)=762.826, p<2.2e-16). That is, within compared to between networks’ edges were larger for direct edges (t(27026)=28.650, p<2.2e-16) and a smaller, opposite effect was found for indirect edges (t(27026)=-8.132, p= p=4.4e-16; SI Fig. 2d). A modest correlation was found between the Euclidian distance and Δ correlation (r(27026)=0.115, p<2.2e-16).

4.4 *Edgewise contribution score across functional networks in DS2*

Examining each network separately, we found a significant positive contribution to structure-function correlation for edges of the visual (t(32)=6.737, p=1.54e-07), control (t(15)=10.178, p=7.50e-08) and dorsal attention (t(14)=2.464, p=0.028) networks and negative contribution for edges in the limbic (t(27)=-3.035, p=0.005) and motor (t(44)=-5.758, p=8.23e-07) networks (SI Fig. 2c). Repeating the test only for edges within the functional networks reveled a significant positive contribution for edges of all networks (all t’s> 2.275, all p’s<0. 023) except for the unaffiliated and the ventral attention networks (t’s<-1.783, all p’s>0.075; SI Fig. 2e).

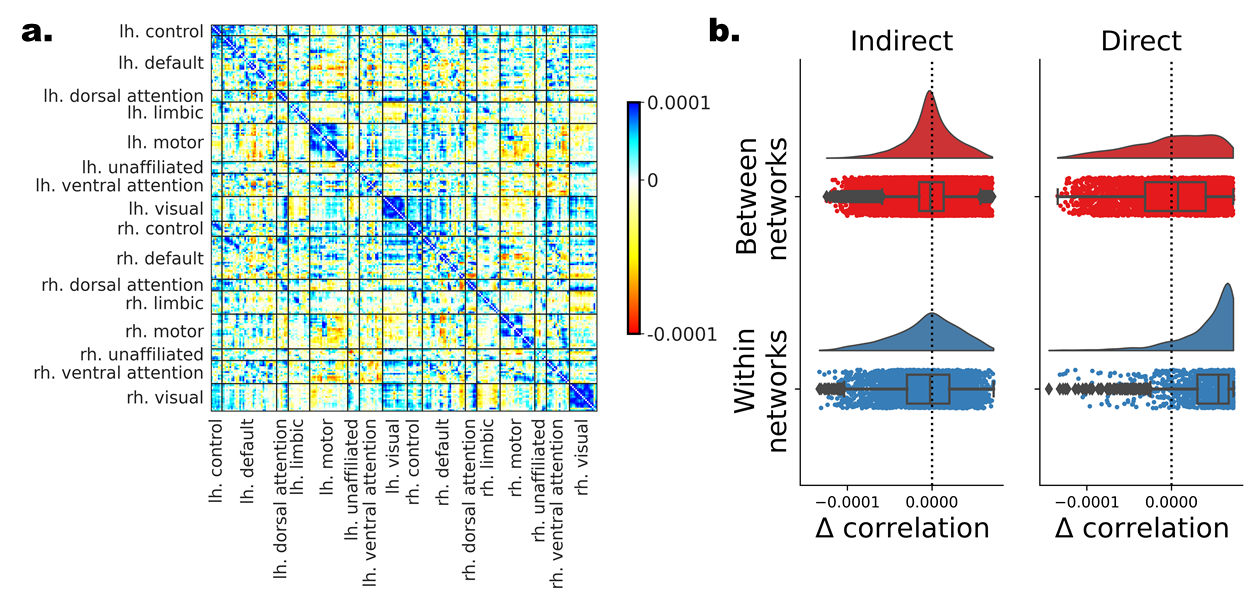

**SI Fig. 2. The contribution of individual edges to the CE-based structure-function mapping in DS2. Individual edges are assessed based on the difference in the measured structure-function correlation after removing each of the edges. (a) All edges depicted in matrix form. Nodes are ordered according to their affiliation to the 7 canonical resting-state networks (Yeo et al., 2011) for the left and right hemisphere separately. (b) Edges with positive contribution to the overall correlation (blue) are those whose elimination results in lower correlation, while edges with negative contribution (red) have the opposite effect. (c) The distribution of Δ correlation for the edges is depicted for all 4 combinations of direct versus indirect and within versus between the 7 networks.**

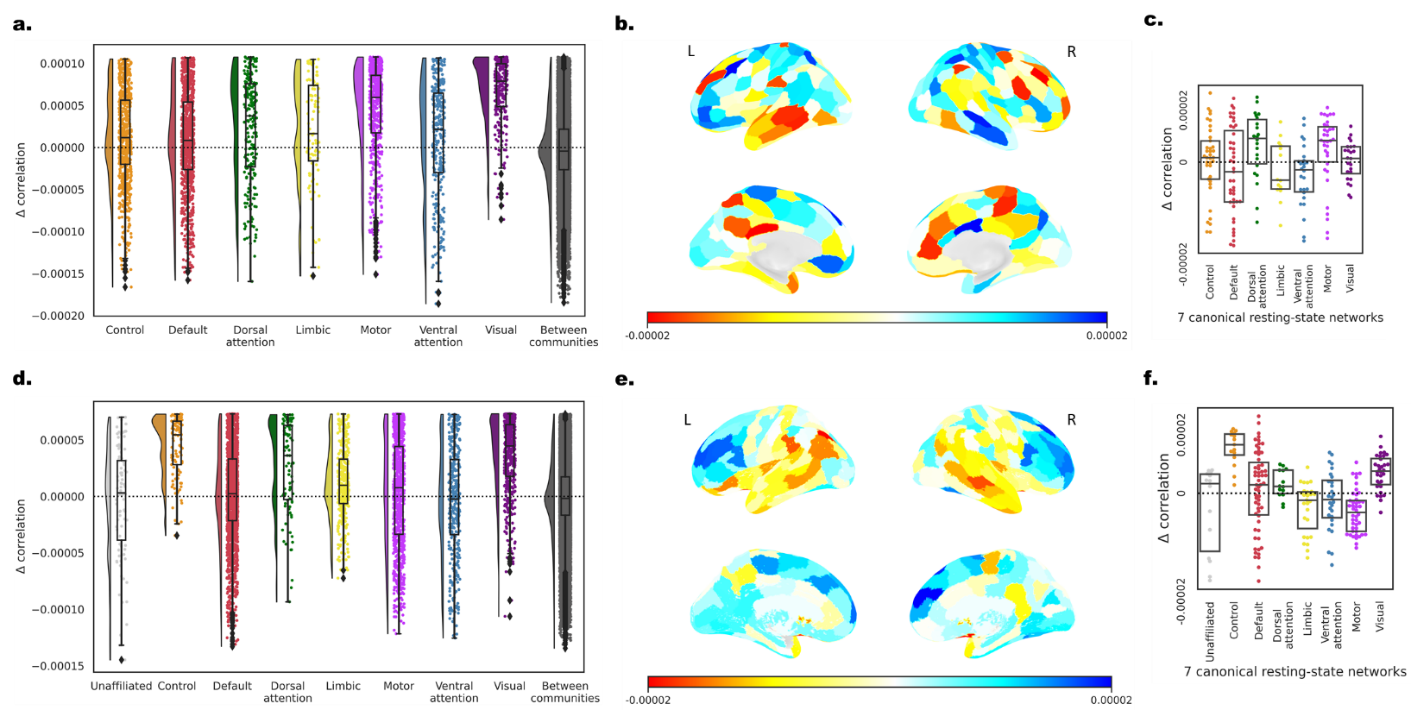

**SI Fig. 3. Contribution of individual edges or nodes to the CE-based structure-function mapping in DS1 and DS2. The distribution of Δ correlation for the edges shown separately within each resting-state network and between communities in (a) DS1 and (d) DS2. Individual edges are assessed based on the difference in the measured structure-function correlation after removing each of the edges. Nodes are ordered according to their affiliation to the 7 canonical resting-state networks (Yeo et al., 2011). (b,e) The average contribution of each node is depicted on the brain surface and (c,f) separately for each of the 7 resting-state networks in DS1 and DS2 respectively. (b,e) The upper row presents a lateral view and the lower row a medial view for both the right (R) and left (L) hemispheres.**

5. *Predicting group-level functional from structural connectivity with deep learning in DS2*

As in DS1, the correlation between the predicted and observed functional connectivity values were ρ(5032)=.542, ρ(21992)=.488 and ρ(27026)=.527; all p’s p<2.2e-16; for the direct, indirect and all edges respectively.

6. *Individual-level structure to function mapping*

The correlation between the direct structural edges and their corresponding functional connectivity edges was significant for all subjects (ρ=.152 ±.028, all p’s<2.2e-16; Fig. S2.d). These correlations were higher when using the CE cosine similarity values (t(144) = 34.4, p<2.2e-16). The mean and standard deviation of the correlation was ρ=.245 ±.040, ρ=.102 ±.035, ρ=.181 ±.344, for the direct, indirect and all edges respectively.

A similar increase in the observed correlation within direct edges was evident both compared to the structural edges (t(144) = 52.8, p<2.2e-16) and the CE cosine similarity measures (t(144) = 34.5, p<2.2e-16; Fig. S3.d). The mean and standard deviation of the predicted-observed FC correlation was ρ=.312 ±.043, ρ=.210 ±.051, ρ=.265 ±.046, all p’s<2.2e-16, for the direct, indirect and all edges respectively.

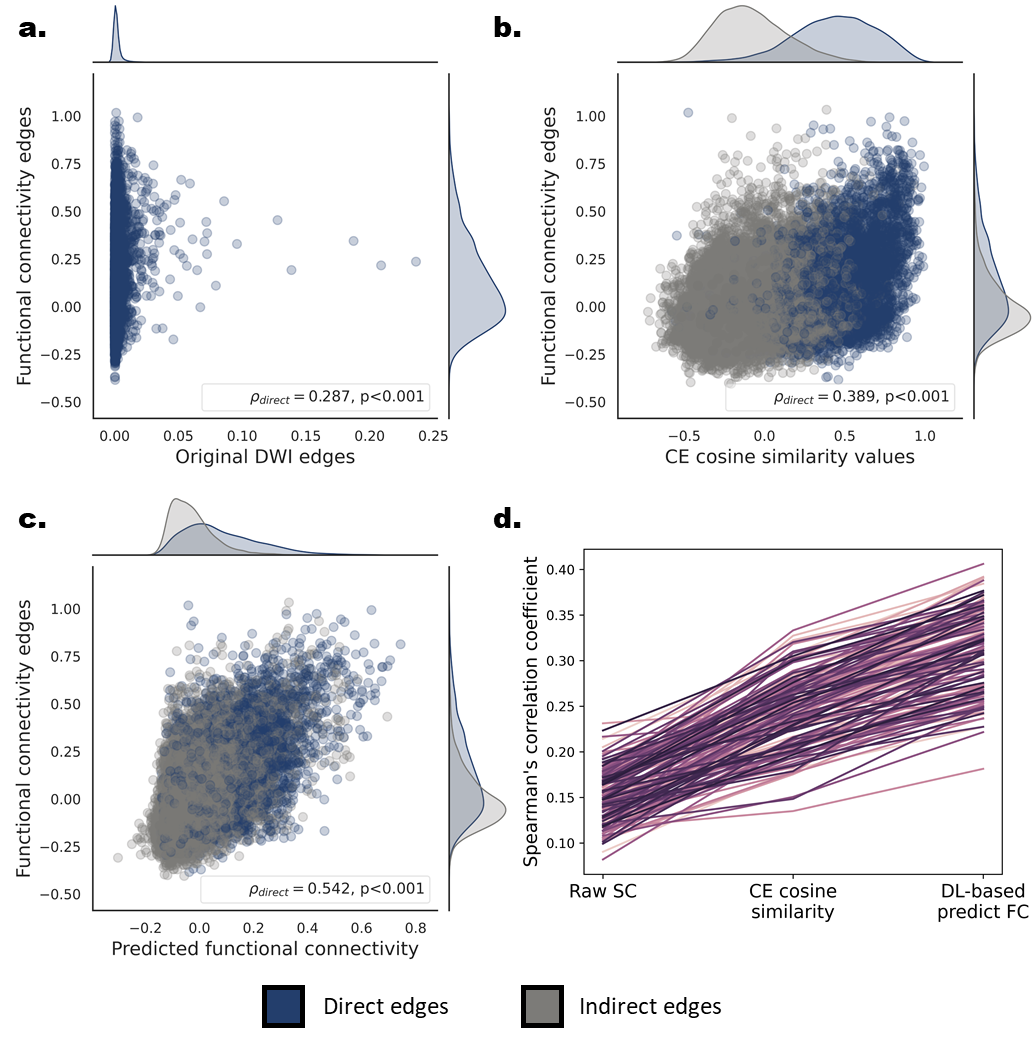

**SI Fig. 4. Correspondence between structural-functional connectivity at the group and individual levels in DS2. (a,b,c) Scatter plots and marginal univariate distributions of functional compared to structural-based edges at the group level. The structure-based edges are the entries of the streamline density matrix (a), the CE cosine similarities (b) and the CE-based predicted functional edges. Direct edges are presented in dark blue and indirect edges in gray. The Spearman’s rank correlations and the corresponding p values for the group-level direct edges are depicted in the bottom right corner of each scatter plot. The correlations for the indirect edges were .256, .488. for the CE cosine similarities and the deep learning based predicted functional edges respectively. (d) Individual-level structure-function Spearman’s correlation values for direct edges of all three structure-based connectivity measures. Each line represents the correlation values of a single subject (n = 145). A significant correlation increase was found for the CE cosine similarities compared to the structural edges and for the CE-based predicted functional edges compared to the CE cosine similarities.**

7. *Parameters of the random walk and their relation to structure to function mapping in DS2*

Shifting from most the locally biased (ρ= 0.202±0.202, 0.002±0.002, 0.049±0.049) to an unbiased random walk (ρ= 0.267±0.267, 0.122±0.122, 0.202±0.202), we observed a significant increase in structure to function mapping (t(11.5)= -12.207, -20.914, -28.666; all p’s<8.76e-12). Moving to the most globally biased random walk (ρ= 0.262±0.262, 0.119±0.119, 0.200±0.200) reveled a smaller, but significant decrease in structure to function mapping (t(11.5)= -2.789, -1.402, -1.267; p=0.010, 0.174, 0.217; see Fig. S3).

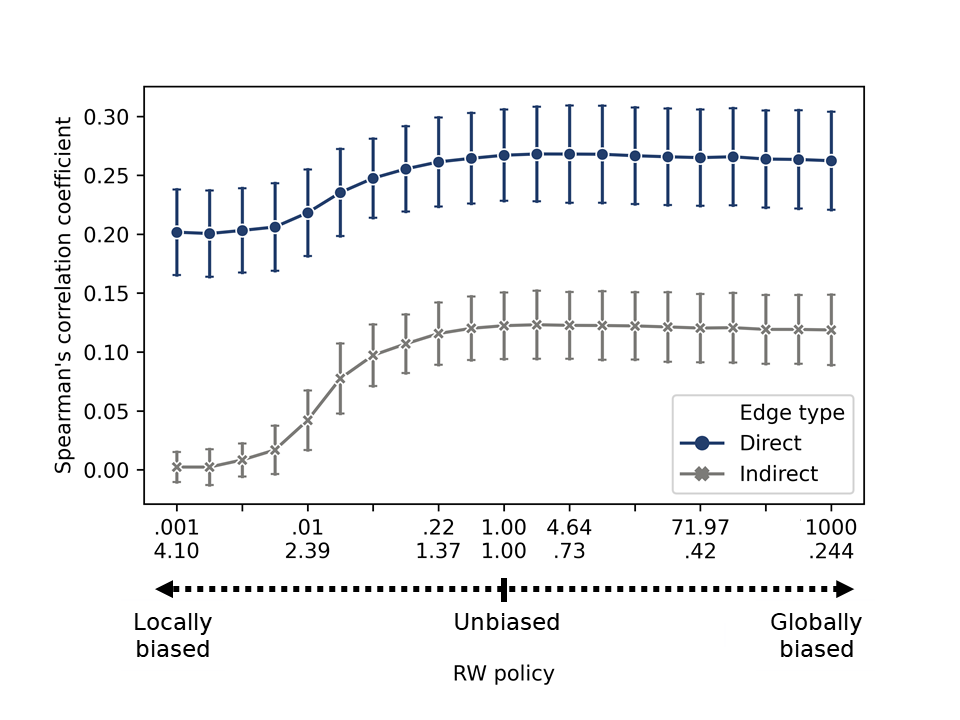
-

**SI Fig. 5. Effect of random walk parameters on structure-function correspondence in DS2. Testing the effect of random walks parameters on the Spearman’s rank correlation between direct (blue) or indirect edges (gray) to their corresponding functional connectivity edges. The random walk parameters were shifted from local (p = 10^-3^, q =4.096) through an unbiased (p = 1, q =1) to global (p = 10^3^, q =0.244) random walk in 20 equal bins on a logarithmic scale. Error bars represent the standard deviation across subjects (N = 25).**

8. *Age-related changes in individual-level structure-function correspondence*

The network contingency analysis method - The network contingency analysis (Sripada et al., 2014) could be used to examined whether each of the canonical resting state network (Yeo et al., 2011) intersections presents a large number of edges correlated with age than expected by chance. This analysis was conducted in the following way: first, age was correlated with each edge across subjects. For each pair of networks intersection or cells of the connectivity matrix (e.g. visual-visual, default-motor), we counted the number of edges whose absolute value was higher than a predetermined threshold. We tested a wide range of thresholds (|r| > 0.25,0.3,0.35,0.4). The number of the suprathreshold edges in each cell was kept and compared to a null distribution created by permuting the age values 10,000 times. A cell was considered to have a significant number of suprathreshold edges if none of the permutations yield a large number than the empirical result. This condition is more conservative than a p<0.1 Bonferroni corrected to the number of tested cells.

Results in DS2 **-** A significant correlation was found between age and the structural edges (r(144)=-.24, p=.003), as in the case of using the CE cosine similarity instead of the SC values (r(144)=-.533, p=5.3e-12). The observed increased correlation was significant (t=4.9, p=1.4e-5). Next, we aimed to identify edges whose contribution to the structure-function correlation increased or decreased with age. We computed the edgewise contribution score for all subjects and correlated each edge with age (Fig. SI5b). Using the network contingency analysis, we revealed significant widespread age-related alteration in the structure-function contribution score (Fig. SI5c). Examining edges within the 7 canonical resting state networks (Yeo et al., 2011), we found a general decrease with age within hemispheres (t’s= 3.83, 2.48, 2.09, 1.19; p’s = 1.3e-4, 0.013, 0.037, 0.233; for |r| threshold of .25, .3, .35 and .4, respectively) and an increase between hemispheres (t’s=14.7, 12.9, 11.1, 9.04; all p’s<2.2e-16; for |r| threshold of .25, .3, .35, .4 respectively). A mix pattern was found between functional networks.

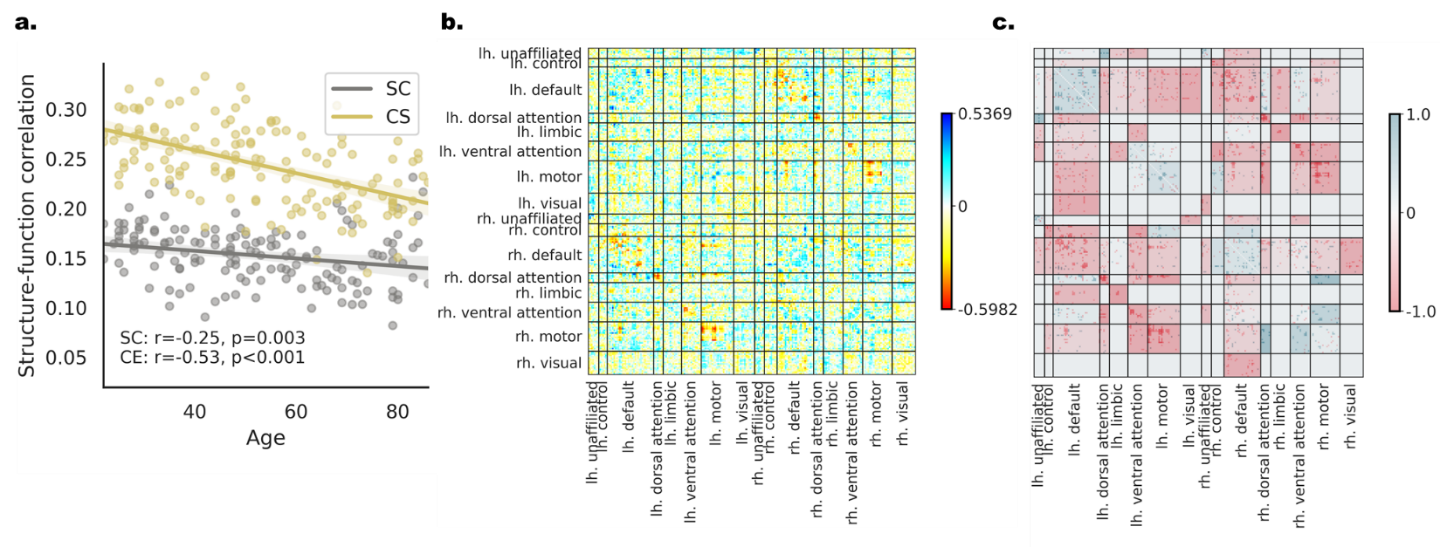

**SI Fig. 6. Age-related changes in individual-level structure-function correspondence in DS2. (a) A regression plot for age and the measured connectome-level structure-function correlation. The values of each subject are presented both for the structural edges (SC, gray) and the CE cosine edges (CE, yellow). The observed correlation to age is indicated on the bottom. The difference in the structure-function correspondence to age is significantly larger for the CE cosine connectivity measure. (b) The correlation of age to the edgewise contribution score of structure-function correspondence is depicted in matrix form. Positive edges are ones that increase the overall correlation with age (blue) and negative edges are ones that decrease the overall correlation with age (red). (c) The result of the network contingency analysis for |r| > 0.25 (see Fig. SI7 for all thresholds) in matrix form. Colored cells had a larger number of edges correlated with age than expected by chance. The cells’ background color is determined by the ratio between edges that are positively (blue) or negatively (red) correlated with age.**

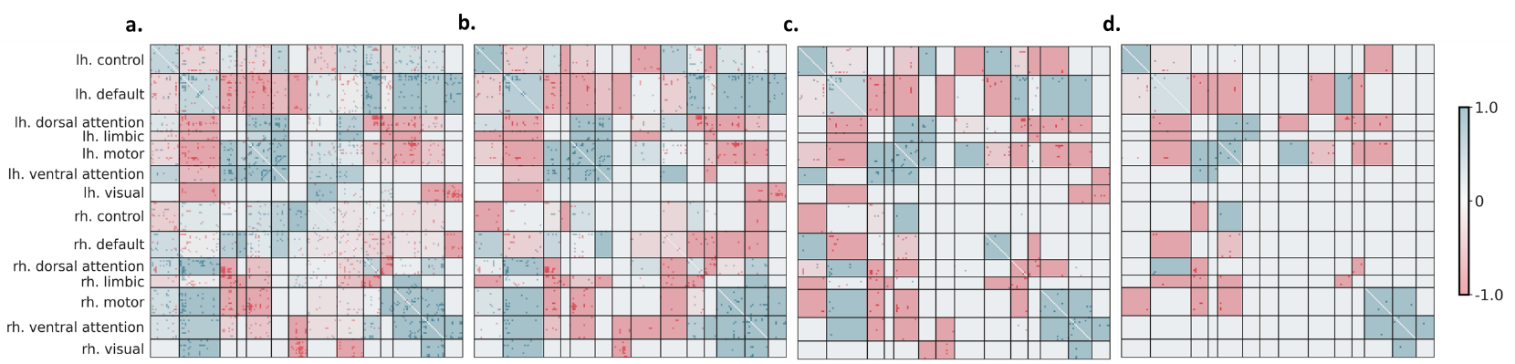

**SI Fig. 7. Network contingency analysis for detecting age-related changes in edgewise structure-function correspondence in DS1. The result of the network contingency analysis for all tested thresholds (|r| > 0.4, 0.35, 0.3, 0.25, for a, b, c, d respectively) in matrix form. Colored cells have a larger number of edges correlated with age than expected by chance. The cells’ background color is determined by the ration between edges that are positively (blue) or negatively (red) correlated with age. Positive edges have larger contribution to the overall correlation with age (blue) and negative edges decrease the overall correlation with age (red).**

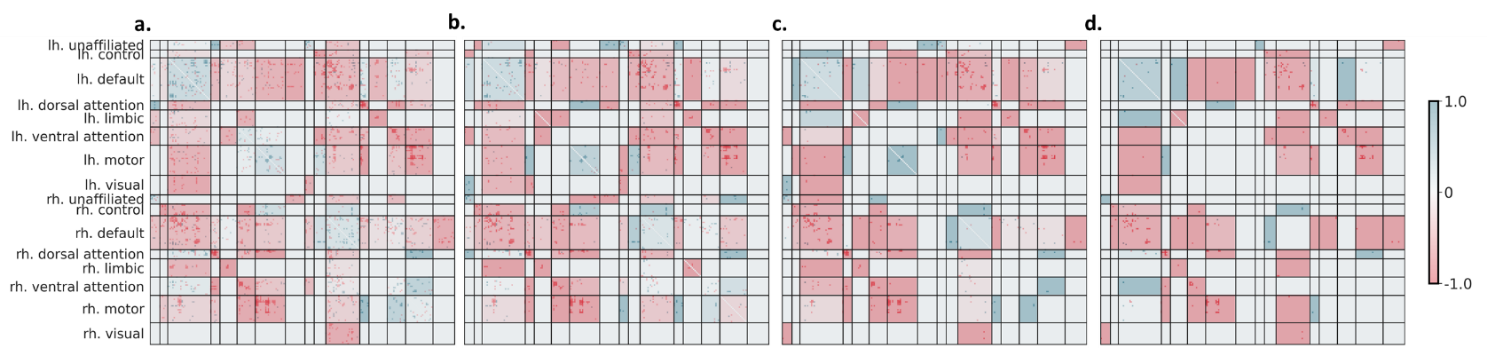

**Fig. 8. Network contingency analysis for detecting age-related changes in edgewise structure-function correspondence in DS2. The result of the network contingency analysis for all tested thresholds (|r| > 0.4, 0.35, 0.3, 0.25, for a, b, c, d respectively) in matrix form. Colored cells have a larger number of edges correlated with age than expected by chance. The cells’ background is determined by the ration between edges that are positively (blue) or negatively (red) correlated with age. Positive edges have larger contribution to the overall correlation with age (blue) and negative edges decrease the overall correlation with age (red).**

9. *Modeling individual differences in age and intelligence using CE*
   9.1 *Capturing individual differences with CE in DS2*

For age, the mean observed-predicted correlation across all nodes was r=.453(±.078) for CE and r=0.367(±0.056) for the CE cosine matrix. And for intelligence r=.240(±.077) for CE, and r=0.138(±0.060) for the CE cosine matrix. The connectome-level model, validated on the left-out test set, revealed a significant correlation of the observed-predicted age (CE: r(199)=0.801, 0.753 , cosine CE: r(199)=0.836, 0.777; for the second-level model and the mean respectively; all p’s<2.2e-16) and intelligence (CE: r(199)=0.578, 0.360, cosine CE: r(199)=0.565, 0.409; for the second-level model and the mean respectively, all p’s<3.5e-07). See SI 9.3 for results after controlling for gender and age.

9.2 *Predictive accuracy of CE compared to structural and functional connectivity in DS2*

The analysis was conducted for a subset of the subjects for whom both functional and structural connectivity data were available (train: 287, test: 138). Using both CE and the CE cosine matrix resulted in significantly better performance than functional and structural connectivity for age (all t’s(232) > 4.36, all p’s <2.0e-05). Similarly, for intelligence (all t’s (232) > 2.53, all p’s <0.012) except for cosine embedding and functional connectivity (t(232) =-0.947p: 0.345). The structural connectivity connectome-level model, validated on the left-out test set, resulted in nonsignificant correlation for both age and intelligence (all r’s(137) < 0.005, p’s >0.415). Contrarily, functional connectivity resulted in significant correlation when taking the mean of the nodal predictions (age: r(137)=0.854 , p=2.2e-16; intelligence: r(137)=0.546, p=4.4e-12) but not for the second-level model (all r’s(137) < 0.039; see SI Fig.9).

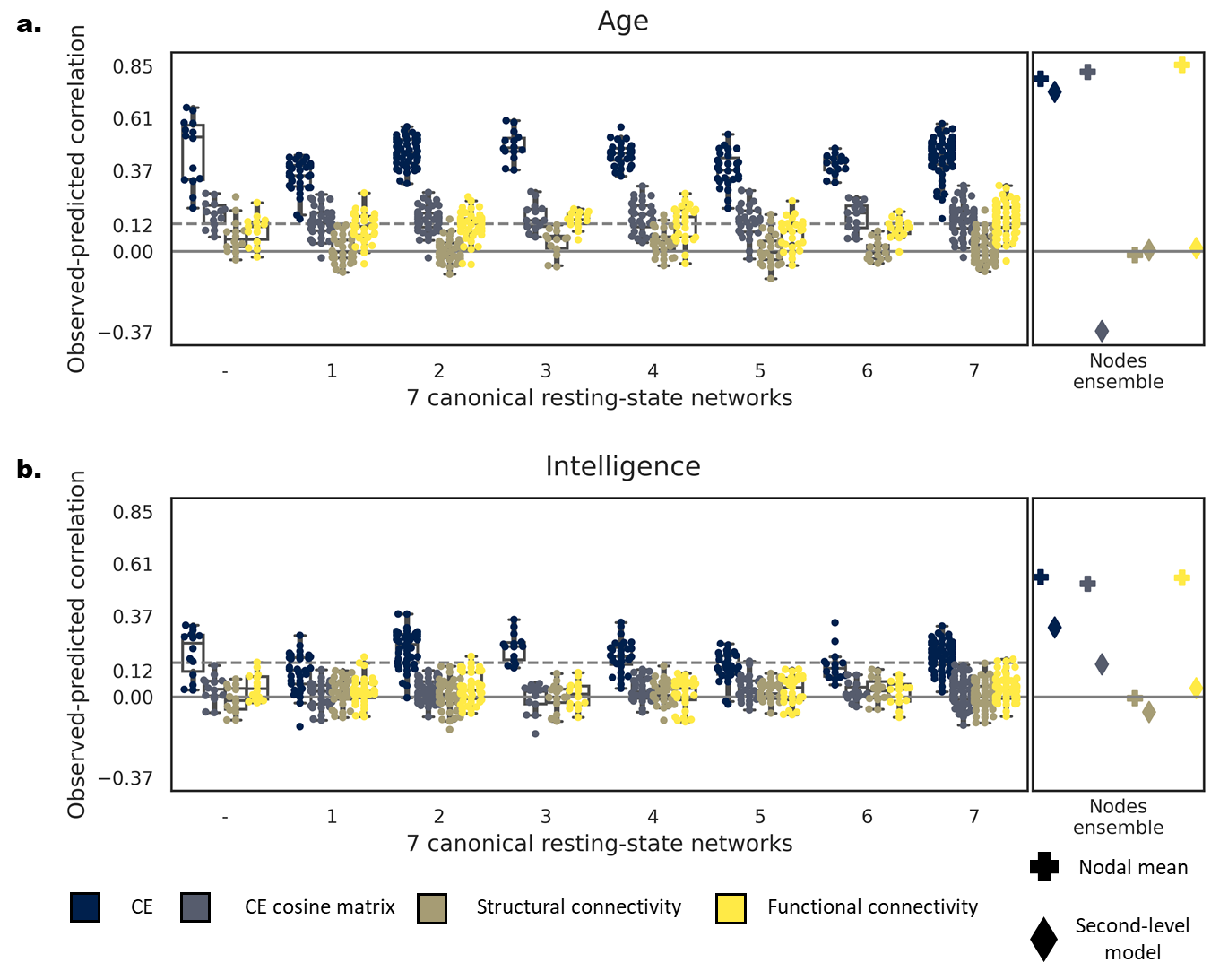

**SI Fig. 9. Comparing predictive accuracy of CE, CE cosine similarity matrix, structural and functional connectivity as input within DS1. Correlation between the observed and predicted age (a) and intelligence (b) with CE (blue), CE cosine matrix (blue-gray), structural (brown) and functional (yellow) connectivity. On the left panel of each subplot, dots represent the predictive accuracy of each node within each of the 7 resting state networks (Yeo et al., 2011). Nodes are grouped by their resting state network affiliation. The right panel depicts the predictive accuracy of a model combining all individual nodes by taking their mean (plus marker) or aggregating them using a second-level linear model (diamond marker). The dashed line represents the FDR-corrected significance level for single nodal prediction.**

9.3 *Capturing individual differences with CE after controlling for age and gender*

We wanted to examine whether our results are confounded by a possible effect of gender, for age prediction, and gender and age, for intelligence prediction. We controlled for the two variables using linear regression by predicting the desired outcome, age and intelligence, with the covariates, gender or age and gender, as predictors, keeping only the residual. **DS1**: For age, the mean observed-predicted correlation across all nodes was r=.448(±.079) for CE and r=0.452(±0.057) for the CE cosine matrix. Similarly, for intelligence r=.083(±.070) for CE, and r=0.074(±0.067) for the CE cosine matrix. The connectome-level model, validated on the left-out test set, revealed a significant correlation of the observed-predicted age (CE: r(188)=0.746, 0.735 , cosine CE: r(188)=0.801, 0.696; for the second-level model and the mean respectively; all p’s<2.2e-16) and intelligence (CE: r(188)=0.384, 0.259, cosine CE: r(188)=0.332, 0.181; for the second-level model and the mean respectively, all p’s<0.016). **DS2**: For age, the mean observed-predicted correlation across all nodes was r=.451(±.078) for CE and r=0.364(±0.056) for the CE cosine matrix. Similarly, for intelligence r=-0.012(±.067) for CE, and r=0.004(±0.067) for the CE cosine matrix. The connectome-level model, validated on the left-out test set, revealed a significant correlation of the observed-predicted age (CE: r(199)=0.798, 0.750 , cosine CE: r(199)=0.833, 0.773; for the second-level model and the mean respectively; all p’s<2.2e-16). For intelligence a small, significant effect was found only with cosine CE second-level model (r(199)=0.150, p=0.038) but not for the other models (all r’s < 0.015, all p’s<0.834).

9.4 *Predictive accuracy of CE compared to structural and functional connectivity after controlling for age and gender*

As in the previous section, we report the results after controlling for gender or age and gender. **DS1**: Using both CE and the CE cosine matrix resulted in significantly better performance than functional and structural connectivity for the age (all t’s(199) > 17.2, p’s <2.2e-16) and intelligence (all t’s (199) > 4.58, all p’s <8.1e-06). The structural connectivity connectome-level model, validated on the left-out test set, resulted in nonsignificant correlation for both age and intelligence (all r’s(188) < 0.119, p’s >0.111). Contrarily, functional connectivity resulted in significant correlation when taking the mean of the nodal predictions (age: r(188)=0.807 , p<2.2e-16; intelligence: r(188)=0.340, p=3.8e-06) but not for the second-level model (all r’s(188) < 0.060). **DS2:** Using both CE and the CE cosine matrix resulted in significantly better performance than functional and structural connectivity for the age (all t’s(137) > 4.24, p’s <3.2e-05) but not for intelligence (all t’s (199)<-0.436). The structural connectivity connectome-level model, validated on the left-out test set, resulted in nonsignificant correlation for both age and intelligence (all r’s(137) < 0.123, p’s >0.150). Contrarily, functional connectivity resulted in a significant correlation when taking the mean of the nodal predictions (age: r(137)=0.852 , p<2.2e-16; intelligence: r(137)=0.179, p=0.036) but not for the second-level model (all r’s(137) < 0.036).
